# Substrate availability and not thermal-acclimation controls microbial temperature sensitivity response to long term warming

**DOI:** 10.1101/2022.09.05.506639

**Authors:** Luiz A. Domeignoz-Horta, Grace Pold, Hailey Erb, David Sebag, Eric Verrecchia, Trent Northen, Katherine Louie, Emiley Eloe-Fadrosh, Christa Pennacchio, Melissa A. Knorr, Serita D. Frey, Jerry M. Melillo, Kristen M. DeAngelis

**Affiliations:** Department of Microbiology, University of Massachusetts, Amherst, MA 01003, USA; Department of Forest Mycology and Plant Pathology, Swedish University of Agricultural Sciences, 750 07 Uppsala, Sweden; IFP Energies Nouvelles, Rueil-Malmaison, France; Institute of Earth Surface Dynamics, Faculty of Geosciences and the Environment, University of Lausanne, Lausanne, 1015, Switzerland; Environmental Genomics and Systems Biology Division, Lawrence Berkeley National Laboratory, Berkeley, CA, United States; The DOE Joint Genome Institute, Lawrence Berkeley National Laboratory, Berkeley, CA, United States; School of Natural Resources and the Environment, University of New Hampshire, Durham, NH 03824, USA; The Ecosystems Center, Marine Biological Laboratories, Woods Hole, MA 02543, USA; Department of Evolutionary Biology and Environmental Studies, University of Zurich, Zurich, 8057, Switzerland

**Keywords:** climate change, soil carbon cycling, carbon use efficiency, microorganisms temperature sensitivity, microbial thermal acclimation

## Abstract

Microbes are responsible for cycling carbon (C) through soils, and the predictions of how soil C stocks change with warming are highly sensitive to the assumptions made about the mechanisms controlling the microbial physiology response to climate warming. Two mechanisms, microbial thermal-acclimation and changes in the quantity and quality of substrates available for microbial metabolism have been suggested to explain the long-term warming impact on microbial physiology. Yet studies disentangling these two mechanisms are lacking. To resolve the drivers of changes in microbial physiology in response to long-term warming, we sampled soils from 13- and 28-year old soil warming experiments in different seasons. We performed short-term laboratory incubations across a range of temperatures to measure the relationship between temperature sensitivity of physiology (growth, respiration, carbon use efficiency and extracellular enzyme activity) and the chemical composition of soil organic matter. We observed apparent thermal acclimation in microbial processes important for C cycling, but only when warming had exacerbated the seasonally-induced, already small soil organic matter pools. Irrespective of warming, greater quantity and quality of soil carbon enhanced the extracellular enzymatic pool and its temperature sensitivity. We suggest that fresh litter input into the system seasonally cancels apparent thermal acclimation of C-cycling processes. Our findings reveal that long-term warming has indirectly affected microbial physiology via reduced C availability in this system, implying that earth system models including these negative feedbacks may be best suited to describe long-term warming impact in soils.

## Introduction

Microbes are key regulators of Earth’s massive soil carbon stocks, determining the partitioning of plant inputs into soil organic matter versus atmospheric carbon dioxide. Climate warming is liable to accelerate microbial activity, changing not only the magnitude but also the composition of soil organic matter stocks. However, the degree to which these changes in microbial activity and soil organic matter will occur is uncertain partly due to insufficient understanding of how the soil-microbe system may acclimate to warming (Bradford et al., 2008; J. M. Melillo et al., 2017; Walker et al., 2018, 2020; Cavicchioli et al., 2019).

Thermal acclimation is the divergence between long- and short-term responses to temperature occurring at the community level and can occur at subcellular to ecosystem scales (Bradford et al., 2008). Observed community thermal acclimation can be the result of both direct and indirect responses to temperature. Direct responses include changes in protein or membrane stability at the cellular level (Hall, Singer, Kainz, & Lennon, 2010; Bradford, 2013) or a shift in the extracellular enzyme pool towards isoenzymes characterized by distinct activation energies and temperature optima (Davidson & Janssens, 2006; Robinson et al., 2017). Indirect drivers of community thermal acclimation can also include changes in resource composition and supply (Kirschbaum, 2004; Moinet et al., 2021; Pold et al., 2016), whose effects on microbial physiology can act through changes in soil microbial community structure (Rocca et al., 2019) and/or reduction of bacterial cellular ribosome content (Söllinger et al., 2022). By shaping the degree to which microbial processes responsible for soil C cycling vary with temperature, thermal acclimation can determine whether soils serve as a source or sink for carbon (Allison, Wallenstein, & Bradford, 2010).

Nonetheless, community thermal acclimation is not a necessary outcome of exposure to long-term elevated temperatures (Carey et al., 2016; Allison, Romero-Olivares, Lu, Taylor, & Treseder, 2018) and can depend on substrate quantity and quality (Alster, Robinson, Arcus, & Schipper, 2022). These substrates change seasonally as plant and microbial inputs fluctuate throughout the year (Wardle et al., 2004; Yarwood, Myrold, & Högberg, 2009; Cheeke, Phillips, Kuhn, Rosling, & Fransson, 2021). Perhaps the most responsive and dynamic of these pools, dissolved organic matter (DOM), consists of compounds derived both directly from root and mycorrhizal exudates and leaf litter leachate, and indirectly from decomposition of soil organic matter (SOM) and the turnover of microbial biomass. Since metabolic rates generally increase with temperature, microbial substrate demand for these substrates should, too (Price & Sowers, 2004). However, not all substrates are equal in their potential to meet microbial demand, whether it be due to differences in enzyme investment required to access them, activation energy needed to be overcome to decompose them, ATP required to uptake them, or cellular reducing equivalents required to assimilate them (Gunina & Kuzyakov, 2022; Gommers, Van Schie, Van Dijken, & Kuenen, 1988). All of these variables affect the carbon use efficiency (CUE), or relative allocation of carbon to growth versus respiration by communities. Notably, the most energetically-rich compounds are not necessarily the most favorable to consume due to the resources which must be invested into extracellular enzymes to access them (LaRowe & Van Cappellen, 2011; Gunina & Kuzyakov, 2022), thereby reducing CUE. Furthermore, SOM components can differ in how sensitive their decomposition is to temperature, leading to considerable changes in soil organic matter chemistry under chronic warming (Pold, Grandy, Melillo, & DeAngelis, 2017; Pec et al., 2021; VandenEnden, Anthony, Frey, & Simpson, 2021; Feng, Simpson, Wilson, Williams, & Simpson, 2008). These long-term warming-induced changes in SOM chemistry are in turn expected to alter the potential resource pool (i.e. DOM) available for organisms to respond to short term changes in temperature (Liu et al., 2021). Since microbial community members differ in their ability to consume various components of DOM (Zhalnina et al., 2018), substrate-dependent responses of microbial physiology to temporary increases in temperature can emerge (S. D. Frey, Lee, Melillo, & Six, 2013). While numerous studies have linked substrate depletion to changes in the apparent temperature sensitivity of microbial respiration (Alster et al., 2022; Bradford et al., 2008; Hartley, Heinemeyer, & Ineson, 2007; Moinet et al., 2021), there is an acute need to understand how potential changes in substrate chemistry under climate warming can affect soil microbial activity. The recent extension of technologies such as Rock-Eval^*®*^ thermal pyrolysis and LC-MS-based metabolomics to soil microbial ecology create opportunities to study how chemistry of soil organic matter and of dissolved organic matter correlate with microbial short-term thermal response, and allow us to begin to disentangle the SOM quality component of apparent thermal acclimation of microbial communities.

Here we used these technologies to test the hypothesis that long-term warming indirectly affects microbial physiology via reductions in the quantity and quality of carbon. We further hypothesised that there is no thermal acclimation of the microbial community to the direct temperature effects of long-term warming, and that apparent differences in microbial physiological thermal response disappear as substrate quantity and quality converge between control and chronically-warmed soils between seasons. We assayed the short-term response of soil microbial physiology following 13 and 28 years of long-term *in situ* soil warming at adjacent field sites within a temperate deciduous forest. Samples were collected from two horizons during mid-summer and autumn in order to reflect natural changes in substrate availability. We used path analysis to assess the interplay between biotic, abiotic, direct and indirect drivers of soil C cycling allowing us to unravel apparent thermal acclimation from changes in soil C available to microbial communities. Our focus was on evaluating the drivers of the temperature sensitivity of respiration and carbon use efficiency, with biotic drivers measured as microbial catabolic potential based on EcoLog assays, extracellular enzyme activity, fungal to bacterial ratios, metagenomics and metabolomics, and abiotic drivers were studied using polar metabolomics to evaluate DOM composition, and ramped-thermal Rock-Eval^*®*^ pyrolysis (RE) to evaluate SOM quantity and quality. Long-term warming did not directly affect microbial physiology according to these analyses, but rather acted indirectly through reductions in the quantity of SOM, supporting our hypothesis.

## Methods

### Soil collection

Soils were collected from two long-term warming experiments at the Harvard Forest Long-Term Ecological Research (LTER) sites in Petersham MA USA (42°30’30”N, 72°12’28”W). The two experiments are located immediately adjacent to each other and had been warmed +5°C for 13 (”SWaN”) (Contosta, Frey, & Cooper, 2011) or 28 (“Prospect Hill” (J. Melillo et al., 2002)) years. Similar disturbance was ensured in heated and control plots at Prospect Hill by installing heating cables in separate disturbance control plots that were never turned on. Experimental plots are 6×6m at Prospect Hill, and 3×3m at SWaN, and follow a randomized block design in the former and a completely randomized design in the latter.

The sites of the two warming experiments are co-located in the same forest stand, characterized by coarse-loamy inceptisols and the same dominant tree species: paper and black birch (*Betula papyrifera* and *lenta*), red maple (*Acer rubrum*), black and red oak (*Quercus velutina* and *rubra*), and American beech (*Fagus grandifolia*). SWaN plots are situated under a canopy gap relative to PH, resulting in slightly different understory vegetation between the two sites (Muth & Bazzaz, 2002). We pooled the control plot data from both sites for analysis since the control plots do not differ in the parameters measured here, with the exception of SOM lability index (I index). Warmed treatment plots at SWaN and PH have not followed the same respiration trajectory over the course of the experiment, so we do not consider the field experiments as two stages in a chronosequence. Additional site details can be found in Supplementary Table S1.

Soils were sampled on July 15^th^ and October 19^th^ 2019, timings which were chosen to reflect soil processes before and after autumn litter deposition (Munger & Wofsy, n.d.). We observed that between July and October of 2019, most of the *Acer* trees had lost their leaves, resulting in an increase in forest floor litter material between the two time points.

Two 10 cm depth soil cores were collected from each of 5 plots per site, treatment, and time point using a 5.7 cm diameter tulip bulb planter. These cores were then separated into organic and mineral soil by colour and texture and sieved to 2 mm on site before being transported back to the lab at ambient temperature, within 4 hours of collection (Figure 1a). We ultimately processed and analyzed a total of 79 samples because the organic soil exceeded the length of the corer and prevented sampling of the mineral soil in one control SWaN plot in July. Immediately upon arrival at the lab, different subsamples of soil were dried to constant mass at 65°C overnight to determine soil moisture content or placed in the −80°C for metabolomic analysis. The remaining soil was left in plastic containers at room temperature (20°C) overnight. Samples which had greater than the target of 48% water holding capacity were left with the lids of the container ajar to allow them to dry slightly overnight (n=2 samples). Soil was then weighed for the CUE, respiration, enzyme assays, microbiological assays and soil organic matter analyses described below (Figure 1).

**Figure. 1.**
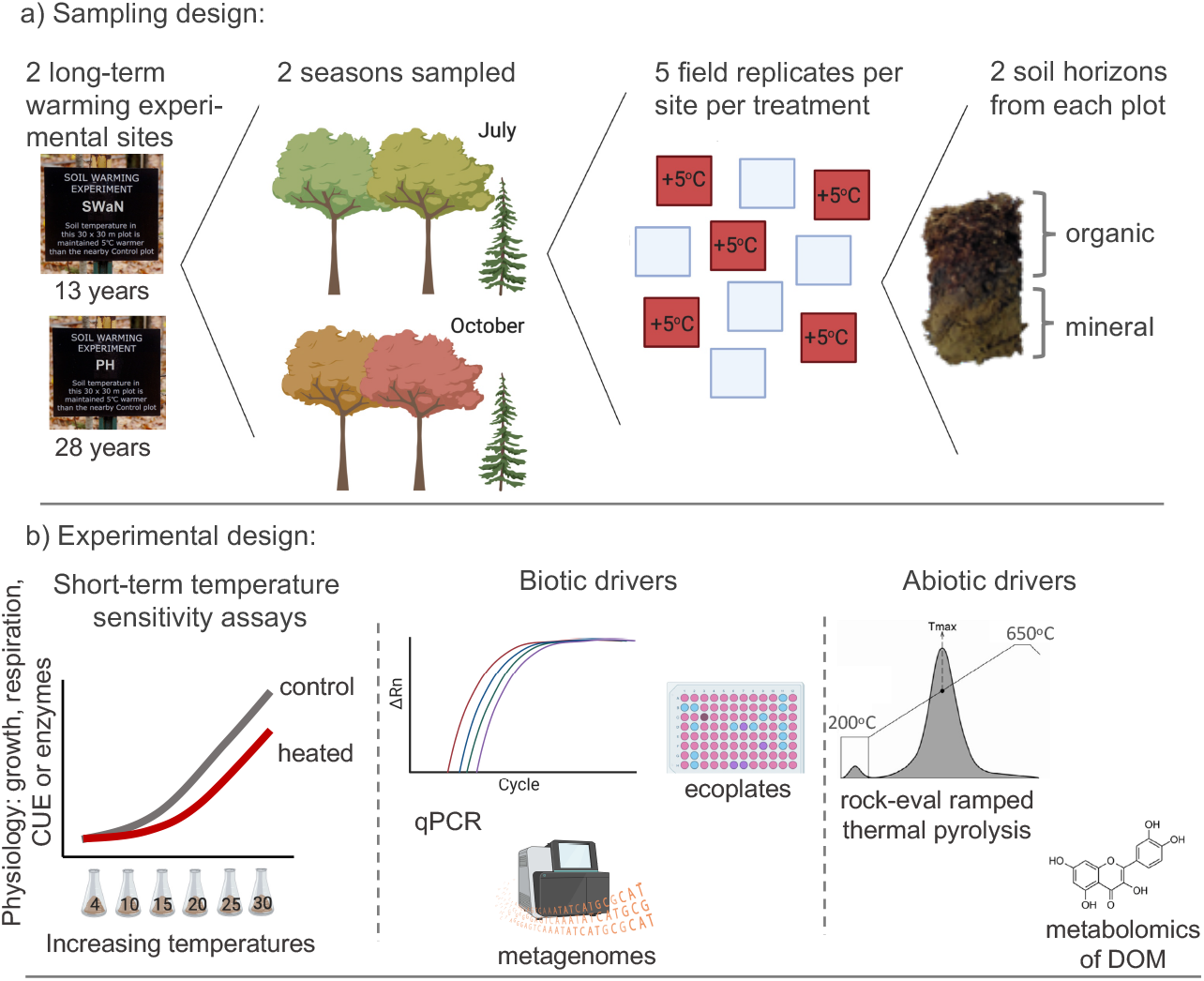
Sampling and experimental design. (a) Sampling design. Samples were collected from two long-term warming experiments, which were established 13 (SWaN) and 28 years (PH) prior to sampling. Samples were collected from the organic and mineral soil horizons in five control and five warmed (5°C above ambient soil temperatures) plots in each experiment, in both July and October. Soils were sieved and transported to the laboratory where we performed a series of assays. (b) Laboratory assays. Temperature sensitivity assays were performed by measuring respiration, growth (^18^O-water method), carbon use efficiency (CUE), and extracellular enzymatic activities (betaglucosidade, N-acetylglucosaminidase, oxidative enzymatic activities between 4 and 30°C). Secondly we also determined the abundance of fungal and bacterial cells with real-time PCR (qPCR), evaluated the potential growth on Biolog EcoPlates at the approximate soil temperature at the time of sampling (e.g. 20 and 25°C for July and 15 and 20°C for October for both heated and control soils); measured the soil water-extractable polar metabolites, and the metagenomes in a subset of data (i.e. organic soils in July). Finally, we measured the soil organic matter quality and quantity using rock-eval ramped thermal pyrolysis and characterized the dissolved organic carbon in soil extracts using LC-MS.

### Microbial community catabolic potential

We used BioLog Ecoplates to evaluate the intrinsic potential growth rate and substrate preferences of microbes in our soil samples. Within 24 hours of collecting soils, we placed 0.25 g (organic) or 0.5 g (mineral) soil in a 125 Erlenmeyer flask with 50 mls of 0.9% sodium chloride and mixed on a shaker table at 180 rpm for 30 minutes. The flasks were then left for five minutes to allow the particles to settle, and then 100 *µ*L of supernatant was transferred to each well of the EcoPlate. The initial absorbance of each plate was measured at 590 and 750 nm and after every 4-12 hours afterwards to capture the full growth curves. The measurements at 750 nm were subtracted from the measurements at 590 nm to isolate the change in dye reduction through time from the change in cell growth. Then the amount of dye colour development in the water only well was subtracted out to quantify the metabolic activity due to the substrate itself. Negative control plates containing only buffer were used to verify the assays were not contaminated. We subsequently fit growth curves to both individual substrates and to the average colour across the plate (mean well colour development) using the *growthcurver* function. Growth rates not significantly different from zero (*P >* 0.05) were set to zero before all substrate-specific growth rates were used for calculating substrate class-specific growth rates and for principal coordinates analysis. Principal coordinates analysis was completed using the vegan package (Oksanen et al., 2019) with a Hellinger-transformed matrix rather than the raw growth rate data so that relative substrate preference could be separated from the overall differences in growth rate under the assay conditions. We refer to the first axis of this ordination as the “ecoplate substrate diversity”.

### Extracellular enzyme activity (EEA)

Between the collection and completion of extracellular enzyme assays (within 4 days), we stored soils at 15oC, which was near the middle of field temperature range. We assayed the cellulose-degrading enzyme *β*-glucosidase (BG), the chitin- and peptidoglycan-degrading enzyme N-acetylglucosaminidase (NAG), and the oxidative enzyme pool (phenol oxidase + peroxidase) at 4, 10, 15, 20, 25 and 30oC, covering most of the range of growing season temperatures experienced by heated and control plot soil communities at these sites. Slurries were prepared with 1.25 g fresh wet weight soil in 175 mls 50 mM pH 4.7 sodium acetate buffer, using a Waring blender set to high for 1 minute. The slurry was stirred at 300 rpm during pipetting to ensure an even distribution of soil. Fourteen technical replicates of 200 *µ*l slurry were pipetted into black plates for BG and NAG, and 500 *µ*l into deep well 2 ml plates for total oxidative enzyme pool. Plates were placed at the assay temperature for at least 25 minutes before substrate addition to allow them to reach the specific target temperature. Fifty *µ*l of 4000 *µ*M 4-Methylumbelliferyl *β*-D-glucopyranoside or 2000 *µ*M 4-Methylumbelliferyl N-acetyl-*β*-D-glucosaminide, or 500 *µ*l 25 mM L-DOPA +0.03% H_2_O_2_ were added to each well, which we previously deemed to be sufficient to attain the maximum activity rate. Each hydrolytic plate contained a slurry-only control (200 *µ*l slurry, 50 *µ*l buffer) and a standard curve (50 *µ*l 0-500 *µ*M in two-fold dilution, in 200 *µ*l buffer). A separate plate for mineral and organic soil quenches (200 *µ*L slurry, 50 *µ*L standard) and substrate-only controls (200 *µ*l buffer, 50 *µ*l substrate) was prepared for each temperature. Hydrolytic assay plates were measured at an excitation/emission wavelength pair of 360/450 nm immediately following substrate addition and after an additional 2 hours. A 100 *µ*l aliquot of the supernatant from oxidative assay plates was removed to a clear polystyrene plate after 4 hours of incubation and read at 460 nm. We used a SpectraMax M2 platereader controlled by the SoftMaxPro v. 5.4 software for all measurements. Extracellular enzyme rates were calculated as previously (German et al., 2011) and normalized by the sample-specific microbial biomass carbon measurement (described below).

### Respiration, growth and CUE

We used the substrate-independent (H_2_^18^O-CUE) method (Spohn, Klaus, Wanek, & Richter, 2016) to evaluate soil microbial carbon use efficiency. Weighing for growth and respiration measurements began the day following soil collection. Three 0.3 g (mineral) or 0.15 g (organic) replicate aliquots of soil were weighed into small vials, placed in a larger tube, and sealed with parafilm to prevent additional moisture loss. Once all samples were weighed (2 days after soil collection), water was added to bring them to 60% water holding capacity. Two replicates received water so that 20% of the total water was present as ^18^O-water, while the remaining replicate received all water as ^16^O-water to account for natural abundance^18^O. Tubes were stoppered with a neoprene bung and immediately placed in the appropriate incubator (4, 10, 15, 20, 25 or 30°C). Empty tubes were also sealed after every 5-10 tubes in order to measure the starting CO_2_ levels in tubes. After 24 (15-30°C) or 48 (4 and 10°C) hours, the CO_2_ was measured in the tube using a 30 ml headspace sample injected into a Quantek instruments infrared gas analyzer with 10 ppm sensitivity. We used a longer time for the lower temperatures because preliminary incubations indicated that it was necessary to reliably detect respiration. The soil samples were placed at −80°C until DNA extraction. We also incubated larger quantities of soil in order to validate respiration measurements from the CUE incubations. 1 g (organic) or 2 g (mineral) soil were brought to 60% water holding capacity with ^16^O-water, and the tubes were incubated alongside the CUE samples.

CUE measurements using the (H_2_^18^O-CUE) method estimate the new microbial biomass produced during the incubation period based on ^18^O-DNA at the end of the incubation period. DNA was extracted from all soils incubated with ^18^O-water and a subset of soils incubated with ^16^O-water using the Qiagen Powersoil kit. Technical duplicates were pooled before quantification using Qubit. The ^18^O enrichment of the DNA was measured using TC/EA-IRMS (Delta V Advantage, Thermo Fisher, Germany) at the UC Davis Stable Isotope Facility. CUE was calculated as per Spohn (Spohn et al., 2016) but using a sample-specific conversion factor rather than the overall average because large differences in MBC:DNA ratios across community types can bias CUE measurements (Pold, Domeignoz-Horta, & DeAngelis, 2020).

### Temperature response curve fitting

We anticipated that respiration and extracellular enzyme activity would increase non-linearly with incubation temperature. Therefore, we compared fits of the Taylor exponential model (Lloyd & Taylor, 1994), and the Ratkowsky square root model (Ratkowsky, Olley, McMeekin, & Ball, 1982) to these data. Models for respiration were fit to individual soil samples using the lm() function in R where the response variable (i.e. respiration or extracellular enzyme activity) was either log- or square root-transformed. We extracted the slope (i.e. temperature sensitivity) estimates from the model for each sample to compare among experimental factors.

Visually, the Ratkowsky model consistently underestimated the respiration and extracellular enzyme activities at higher temperatures compared to the Taylor model, and the R^2^ statistic was also consistently lower (median=0.052). Therefore, we proceeded to extract the slope parameter from the Taylor model and used this as a metric of temperature responsiveness for downstream statistical analyses. The growth and CUE data were very variable compared to the respiration data and individual samples did not follow a consistent pattern. Therefore, we could not fit a curve to the data and instead used ranked CUE and growth rates at different temperatures for each soil sample to determine the optimum temperature and whether it differed between warming treatments.

### Quantitative PCR (qPCR)

The abundance of total fungi and total bacteria was assessed using qPCR with ITS primers (ITS1: 5’-TCCGTAGGTGAACCTGCGG-3’ and 5.8S: 5’-CGCTGCGTTC TTCATCG-3’) and 16S ribosomal RNA primers (Eub338: 5’-ACTCCTACGGGAGGCAGCAG-3’ and Eub518: 5’ATTACCGCGGCTGCTGG-3’) (Fierer, Jackson, Vilgalys, & Jackson, 2005), respectively. The abundance in each soil sample was based on increasing fluorescence intensity of the SYBR Green dye during amplification. The qPCR assay was carried out in a 15 *µ*l reaction volume containing 2 ng of DNA, 7.5 *µ*l of SYBR Green PCR master mix (Qiiagen quantifast SYBR kit) and each primer at a concentration of 1 *µ*M. Inhibition tests were performed by running serial dilutions of DNA extractions and did not indicate inhibition of amplification. For each sample at least two independent qPCR assays were performed for each gene with technical duplicates within each assay. The qPCR efficiencies for both genes ranged between 78 and 110%. Values are reported as gene copy number g^-1^ dry soil.

### Microbial biomass carbon (MBC)

Quadruplicate 0.5g (organic) or 1g (mineral) soil samples for microbial biomass carbon (MBC) were weighed out with the CUE samples. We kept the weighed out soil samples at 15°C (the middle of the temperature range used for CUE and enzyme measurements) until the end of CUE incubations so growth and biomass measurements could be taken on similarly-treated soils. Two replicates were exposed to chloroform vapor fumigation under vacuum for 24 hours, while the other two replicates were placed at 4°C for the duration. Soils with and without chloroform were subsequently extracted in 15 mls of 0.05M K_2_SO_4_, which a preliminary trial indicated led to similar levels of extractable organic carbon and MBC detected as the more standard 0.5 M K_2_SO_4_ for our soils. DOC was measured on a Shimadzu TOC analyzer.

### Soil organic matter quality

We used Ramped-thermal Rock-Eval^*®*^pyrolysis (RE) to evaluate SOM quality (Soucémarianadin et al., 2018). During RE, carbon oxides are quantified as they come off a soil sample subject to increasing temperatures, thereby providing a metric of SOM intrinsic thermal stability. Compounds with high thermal stability include aromatic and phenolic non-lignin compounds, while lipids and polysaccharides tend to have lower thermal stability (Sanderman & Grandy, 2020). Mineral and organic soils were dried at 65°C and crushed to a fine powder in a mortar and pestle. Between 50-70 mg soil was pyrolyzed over a temperature ramp from 200 to 650oC, followed by combustion to 850°C using a Rock-Eval 6 pyrolyser (Vinci technologies) at the Institute of Earth Sciences of the University of Lausanne (Switzerland). Hydrocarbons released during this process were measured by a flame ionization detector (FID). The resultant thermogram was used to calculate the I index (”labile carbon fraction”) and R index (“recalcitrant carbon fraction”) as previously (Sebag et al., 2016). We also used the thermogram of each sample to calculate a Bray-Curtis distance matrix of all samples, taking the peak height at each 1°C increment as an input value to create the distance matrix between all samples (Domeignoz-Horta et al., 2021). This approach allowed us to evaluate SOM composition in addition to C quantities in the different soils. The first axis of the NMDS was used as a proxy for SOM quality as it was strongly correlated with the I index (R^2^=0.89, *P* ¡ 0.0001).

### Metabolomics

Water extractable polar metabolites (here, “water-extractable organic matter”, or WEOM) were obtained by shaking 1 g dry soil equivalent with 5 ml (mineral soil) or 10 ml (organic soil) of LC/MS-grade water at 200 RPM and 4°C for 1 hour. Every extraction was accompanied with a water-only control. The soil extracts and the water controls were filtered through a 45 *µ*m PTFE filter. The filtrate was lyophilized and sent for analysis at the DOE Joint Genome User Facility at Lawrence Berkeley National Laboratory. Polar metabolites were resuspended in 170 *µ*L of 100% methanol containing 13C-15N labeled amino acids (30 *µ*M, 767964, Sigma). UHPLC normal phase chromatography was performed using an Agilent 1290 LC stack, with MS and MS/MS data collected using a Q Exactive HF Orbitrap MS (Thermo Scientific, San Jose, CA). Full MS spectra was collected from m/z 70 to 1050 at 60k resolution in both positive and negative ionization mode, with MS/MS fragmentation data acquired using stepped 10, 20 and 40 eV collision energies at 17,500 resolution. Mass spectrometer source settings included a sheath gas flow rate of 55 (au), auxiliary gas flow of 20 (au), sweep gas flow of 2 (au), spray voltage of 3 kV and capillary temperature of 400 oC. Normal phase chromatography was performed using a HILIC column (InfinityLab Poroshell 120 HILIC-Z, 2.1 150 mm, 2.7 *µ*m, Agilent, 683775-924) at a flow rate of 0.45 mL/min with a 2 *µ*L injection volume. To detect metabolites, samples were run on the column at 40 °C equilibrated with 100% buffer B (99.8% 95:5 v/v ACN:H_2_O and 0.2% acetic acid, w/ 5 Mm ammonium acetate) for 1 minute, diluting buffer B down to 89% with buffer A (99.8% H_2_O and 0.2% acetic acid, w/ 5 mM ammonium acetate and 5 *µ*M methylene-di-phosphonic acid) over 10 minutes, down to 70% over 4.5 minutes, down to 20% over 0.5 minutes, and isocratic elution for 2.25 minutes, followed by column re-equilibration by returning to 100% B over 0.1 minute and isocratic elution for 3.9 minutes. The injection order of the 79 experimental samples and 5 extraction controls was randomized and an injection blank (2 *µ*L of 100% MeOH) run between each sample. Metabolomics data were analyzed using Metabolite Atlas (Yao et al., 2015) and with in-house Python scripts to obtain extracted ion chromatograms and peak heights for each metabolite (Supplementary table S2). Metabolite identifications were verified with authentic chemical standards and validated based on three metrics comparing (1) detected versus theoretical m/z (*<* 5ppm), (2) retention time (≤ 0.5 min) and (3) fragmentation spectra similarity to a chemical standard run using the same chromatography and LC-MS/MS method. Data from internal standards and quality control samples (included throughout the run) were analyzed to ensure consistent peak heights and retention times. Peak height of different samples were calculated out of 4 replicates and we excluded metabolites that showed a quality score under 1 and were not measured in every sample. Where multiple ions existed for the same compound within a sample analyses we summed the peak heights. Where compounds appeared in both the positive and the negatively charged compound summary table, we used the greatest value. After quality control we had 227 compounds. The WEOM matrix was normalized by sum and pareto scaled before calculating Bray-Curtis distance and completing principal coordinates analysis using the *MetabolAnalyze, vegan* and *ape* packages respectively (Nyamundanda, Brennan, & Gormley, 2010; Oksanen et al., 2019; Paradis & Schliep, 2019). Pareto scaling divides each compound by the square root of the variance. We selected this scaling approach to moderately reduce the importance of the extremely dominant compounds and remove the large differences in total peak height between mineral and organic soil samples without completely abolishing all relative abundance information.

Compound complexity was determined based on the Bertz/Hendrickson/Ihlenfeldt complexity reported in PubChem. The complexity score is higher in polymers compared to monomers of the same compound, and in asymmetric and/or heteropolymeric compared to symmetric and/or homopolymeric compounds. Therefore, the complexity score can be considered an imprecise proxy for how difficult a compound is to break down. These values were downloaded from PubChem. Mean nominal oxidation state was calculated as per (LaRowe & Van Cappellen, 2011) (NOSC=-((-Z + 4C + H - 3N - 2O + 5P −2S)/C) + 4). NOSC was selected as a proxy for the quality of DOC because oxidation of compounds with higher NOSCs is associated with greater electron transfer (LaRowe & Van Cappellen, 2011), while compounds with lower NOSCs have greater potential energy but require a larger investment to acquire it. As such, the NOSC of SOM generally decreases as decomposition proceeds (Gunina & Kuzyakov, 2022). The NOSC and complexity indices for each compound are reported in Supplementary Table 3.

### Metagenome sequencing and annotation

Metagenomes were obtained from organic soil collected in July from both sites, totaling 20 samples. DNA extracted from soils incubated at 20°C was sequenced at the Joint Genome Institute (Supplementary methods and supplementary table 2). Reads were assembled and PFAMs were annotated in the assembled fraction of reads using the DOE-JGI Metagenome Annotation Pipeline (Huntemann et al., 2016). This led to an average of 1.72 Gbp assembled and 2,807,085 genes predicted per metagenome. We then mapped PFAMs to CAZys (Pold et al., 2016) and used mean mapping depth for CAZys relative to the mapping depth to the single copy *rpoB* gene (PF04563, PF04561, PF04565, PF10385, PF00562, PF04560) as our proxy of CAZy abundance across samples. We elected to focus on CAZymes as they are responsible for the polymerization and depolymerization of many of the dominant compounds found in leaf litter and microbial cell walls, including cellulose, lignin, and chitin. We subsequently categorized CAZymes based on the primary substrates they decompose (Berlemont, Martiny, & Kivisaar, 2015).

### Data Analysis

Analysis was completed in R version 3.6.3 using packages reshape2 (Wickham, 2007), vegan (Oksanen et al., 2019), lme4 (Pinheiro, Bates, DebRoy, Sarkar, & R Core Team, 2019), agricolae (Mendiburu, 2019), grofit (Kahm, Hasenbrink, Lichtenberg-Frat’e, Ludwig, & Kschischo, 2010), dplyr (Wickham, François, Henry, & Müller, 2022), lattice (Sarkar, 2008), MASS (Venables & Ripley, 2002) and permute (Simpson, 2022). Variables were log-transformed where necessary to fit model assumptions. We calculated Hedge’s G (Cohen, 1988) using the effect size package (Torchiano, 2020) to compare the effects of warming treatment and season on biotic and abiotic soil properties; we considered drivers to have a significant effect when the 90% confidence intervals surrounding the effect size estimate did not overlap zero (Figure 2). We elected to use an alpha of 0.1 (and 90% confidence intervals) rather than 0.05 (or 95% confidence intervals) because we were limited by the number of field replicates we could collect and a power analysis using means for microbial biomass, soil C stocks, and extracellular enzyme data collected from the site in 2014 (Pold et al., 2017) indicated this would be needed to detect changes in mean values of 25% with a power of 80%. Figures were made using the ggplot2 package (Wickham, 2016).

**Figure 2.**
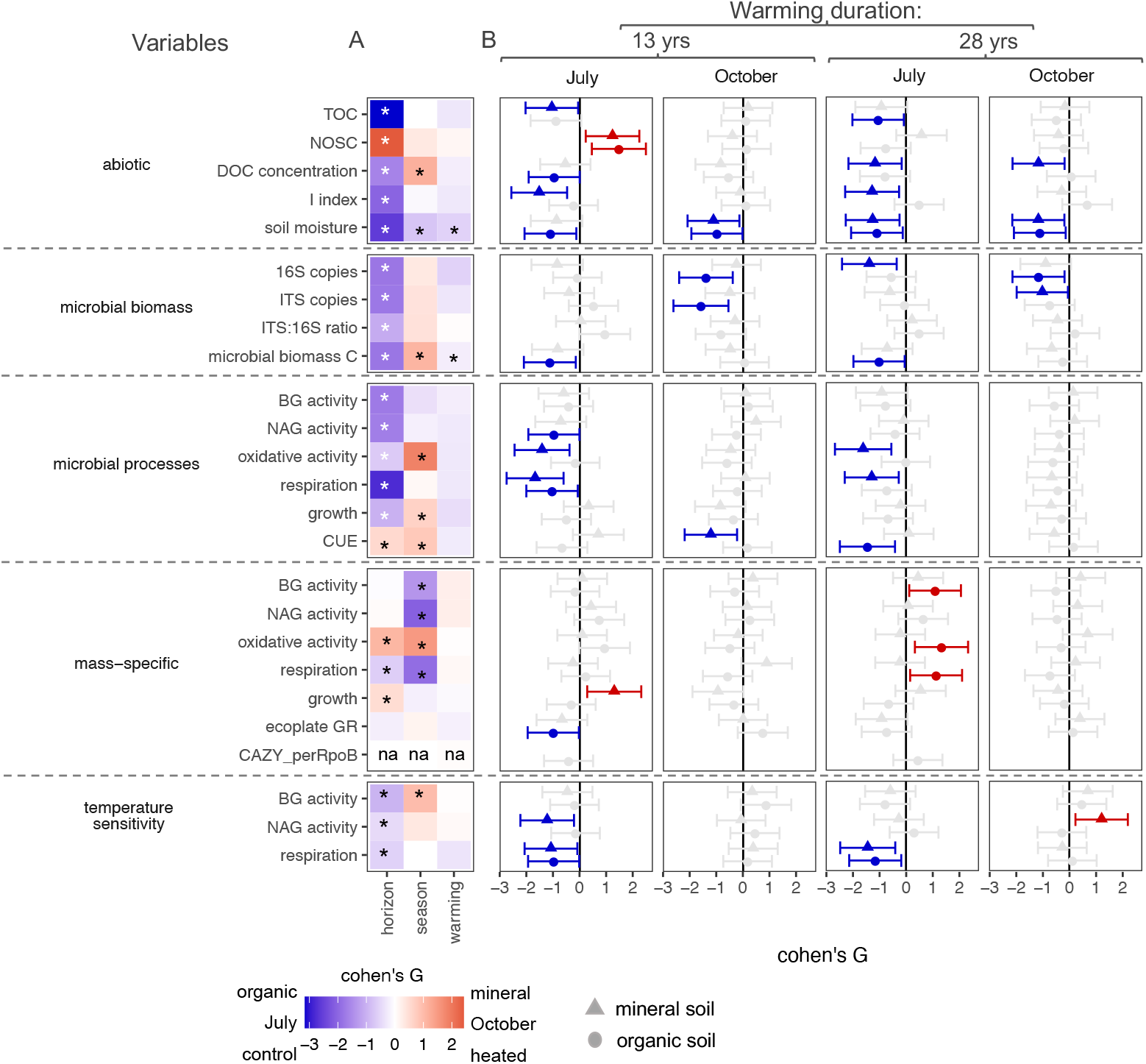
Effect of experimental variables on abiotic conditions and microbial physiology. The variables surveyed were soil properties, measurements of microbial biomass, microbial activity, mass-specific microbial activity, and the temperature sensitivity of microbial activity. The heatmap shows overall differences between horizons, sampling seasons, and warming treatments (A), while the effect size plots show the effect of warming treatment within a warming experiment and sampling season (B). Both are based on Cohen’s G, with asterisks in A and red (heated > control) and blue (heated < control) shapes in B denoting instances where the 90% confidence intervals do not overlap zero. Shapes in B differentiate mineral (triangles) from organic (circles) soils.

### Structural equation modeling

We used structural equation modeling (SEM) to test the hypothesis that long term warming affects microbial carbon use efficiency and temperature sensitivity of respiration via changes in C quantity and quality. The hypothesized path structure for the temperature sensitivity of respiration was based on the premise that long-term warming, season and soil depth had direct effects on C quantity, quality and microbial communities (fungal:bacterial ratio). These variables are then hypothesized to affect the size and temperature sensitivity of the extra-cellular enzyme pool and the composition of the water-extractable soil organic matter. Finally, we hypothesized that the temperature sensitivity of respiration would be controlled by the enzyme pool, the enzyme temperature sensitivity and the quantity and quality of soil C (Supplementary Figure S10). We also completed SEM analysis on microbial CUE at 20°C. This model had a similar overall layout of the temperature sensitivity of respiration model, except that we additionally hypothesized that diversity of EcoPlate substrate consumption would be affected by season and carbon quality, and that respiration and growth would be impacted by carbon quantity and quality (Supplementary figure S11).

The SEM model path fit was performed using the piecewiseSEM package (Lefcheck, 2016). Cumulative enzyme activity and temperature sensitivity are represented in the model as composite variables (Lefcheck, 2016), using the mean of BG, NAG and oxidative enzymes for the former and just of the hydrolytic enzymes for the latter. We kept the model that explained the most variation in temperature sensitivity of respiration and CUE, and had a non-significant Chi-squared test (*P >* 0.05), low Akaike Information Criterion (AIC), and high Comparative Fit Index (CFI *>* 0.9). If the test of direct separation (Lefcheck, 2016) identified missing paths in our hypothesized model we added these paths into the model.

## Results

### Long-term warming affects soil C substrates

Warming reduced soil organic matter quality and quantity by most measures, but this effect was principally visible during the July sampling (Figure 2). We evaluated the chemistry of soil dissolved organic matter by analyzing the polar water-extractable fraction of soil organic matter (WEOM) using metabolomics, and the thermostability of soil organic matter using Rock-Eval^*®*^.

The nominal oxidation state of compounds (NOSC) identified in the WEOM varied from −1.5 to 3, though the mean value was less than zero for all samples (Supplementary Figure S7). The effect of warming on NOSC was site and time point dependent, increasing only with warming in SWaN plots in July (Figure 2, Supplementary Figure S7). Warming and sampling time point did not affect the molecular complexity of the WEOM, but molecular complexity was higher in mineral compared to organic soil (Figure 2a, Supplementary Figure S7, mean of 143 vs. 135). Overall dissimilarity in WEOM composition was primarily explained by horizon (Adonis R^2^=0.46; ADO-NIS F(1,78)=87.50, *P <* 0.001), followed by warming treatment (Adonis R^2^=0.083; F(2,78)=7.82, *P <* 0.001) and time point (Adonis R^2^=0.045; F(1,78)=8.46, *P <* 0.001). There were no significant interactions between these variables, and no evidence of heterogeneous group dispersions (i.e. betadisper *P >* 0.05).

Long-term warming reduced the total dissolved organic carbon in the mineral soil at the 28 year-old warming experiment at both timepoints, and reduced the lability index of the total soil organic matter pool in July in both field experiments (Figure 2). Warming explained just 2.2% of the variation in SOM chemical composition based on ramped thermal Rock-Eval^*®*^ (F(1,76) = 7.30, *P* = 0.006), with much of the remaining variation explained by soil horizon (Adonis R^2^=0.71, F(1,76) = 231.22, *P <* 0.001) and, to a lesser degree, season (R^2^=0.020, F(1,76) = 6.76, *P* = 0.008)(Figure 2). However, the effect of warming depended on horizon (warming x horizon interaction R^2^=0.011, F(1,76) = 3.76, *P* = 0.04) and sampling timepoint (sampling x warming interaction R^2^=0.008, F(1,76) = 2.70, *P* = 0.09). The first axis of this ordination was strongly positively correlated with the SOM I index (inset, Figure 4a, cor: 0.89 *P <* 0.001).

### Long-term warming affects microbial communities and activity

Effects of warming were most apparent in July (Figure 2). Microbial biomass carbon was reduced by up to 45% in July (Figure 2a and Supplementary Figure 1). Warming also reduced the July respiration at 20°C in mineral soils at both field experiments, in addition to reducing respiration in organic soil samples taken from the 13-year old field experiment (Figure 2). Select extracelullar enzyme activities per gram of soil were also reduced by warming in both sites in July, but oxidative and BG enzyme activities per unit microbial biomass were enhanced by warming in organic soils of the older field experiment (Figure 2b). Based on cultivation in EcoPlates, potential growth rate of microbial communities was unaffected by warming, except for the organic horizon of the 13 year-old site in July (Figure 2b).

To understand how microbial communities might adapt to long-term warming in these soils we measured the response of some microbial processes to temperature from 0 to 30°C. Expected increases in process rates with the incubation temperatures (4-30°C) could only be detected for respiration and extracellular enzyme activity (Figure 3a-d; Supplementary Figure S5). The temperature sensitivity of respiration was reduced by long-term warming in July in both long-term warming experiments and soil horizons (Figure 2 and 3a). While BG activity showed a trend of smaller activity in heated compared to control soils, this difference was not significant (Figure 2 and Figure 3d), but NAG showed a lower response to temperature in heated soils in July within the 13-years site (Figure 2 and Supplementary Figure S6).

**Figure 3.**
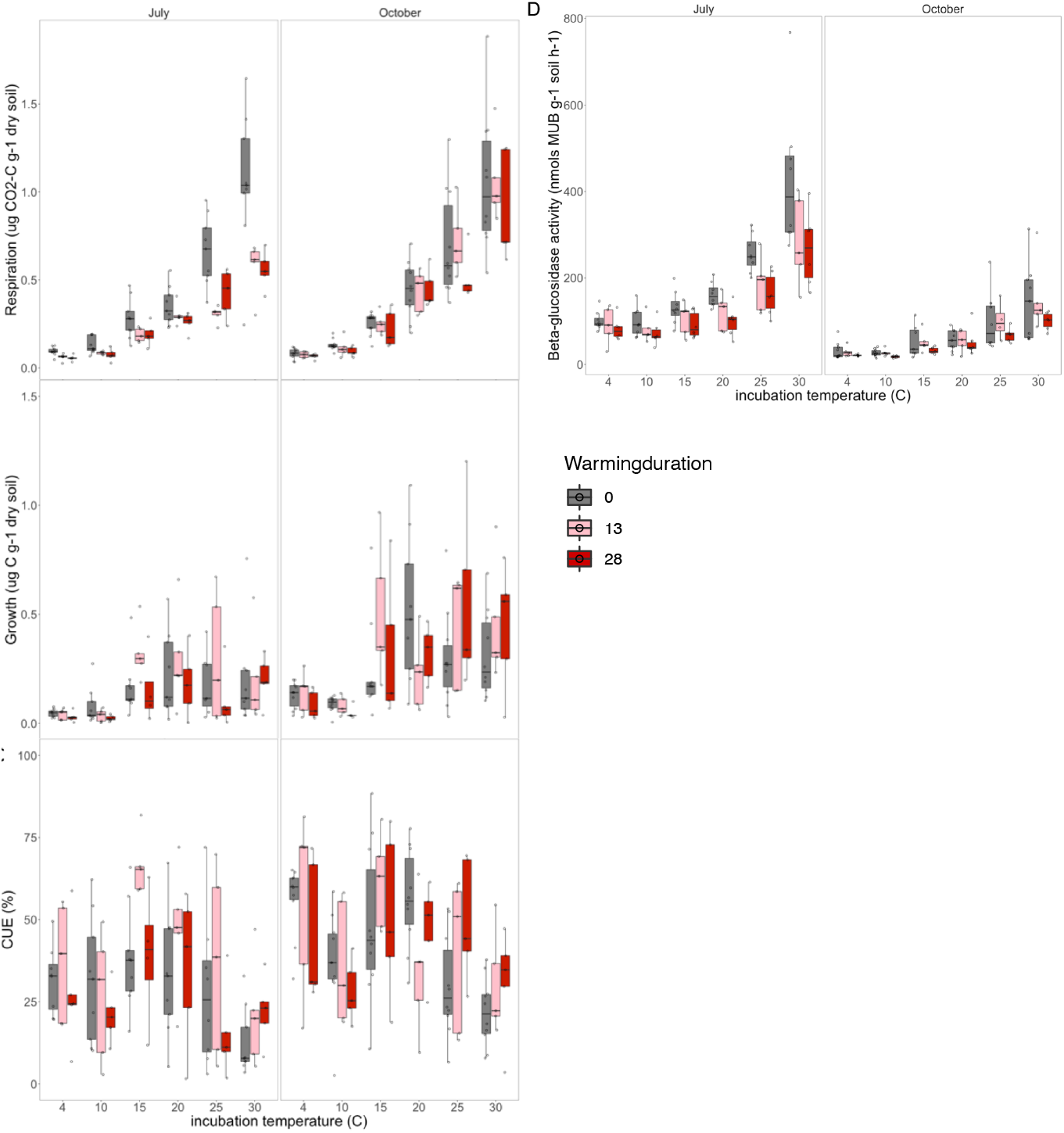
Boxplots of temperature sensitivity of microbial carbon cycling processes. Respiration rate (A), growth rate (B), CUE (C) and BG extracellular enzyme activity (D) at different temperatures from 4 to 30°C measured under laboratory incubations in July and October for the mineral soils.

**Figure 4.**
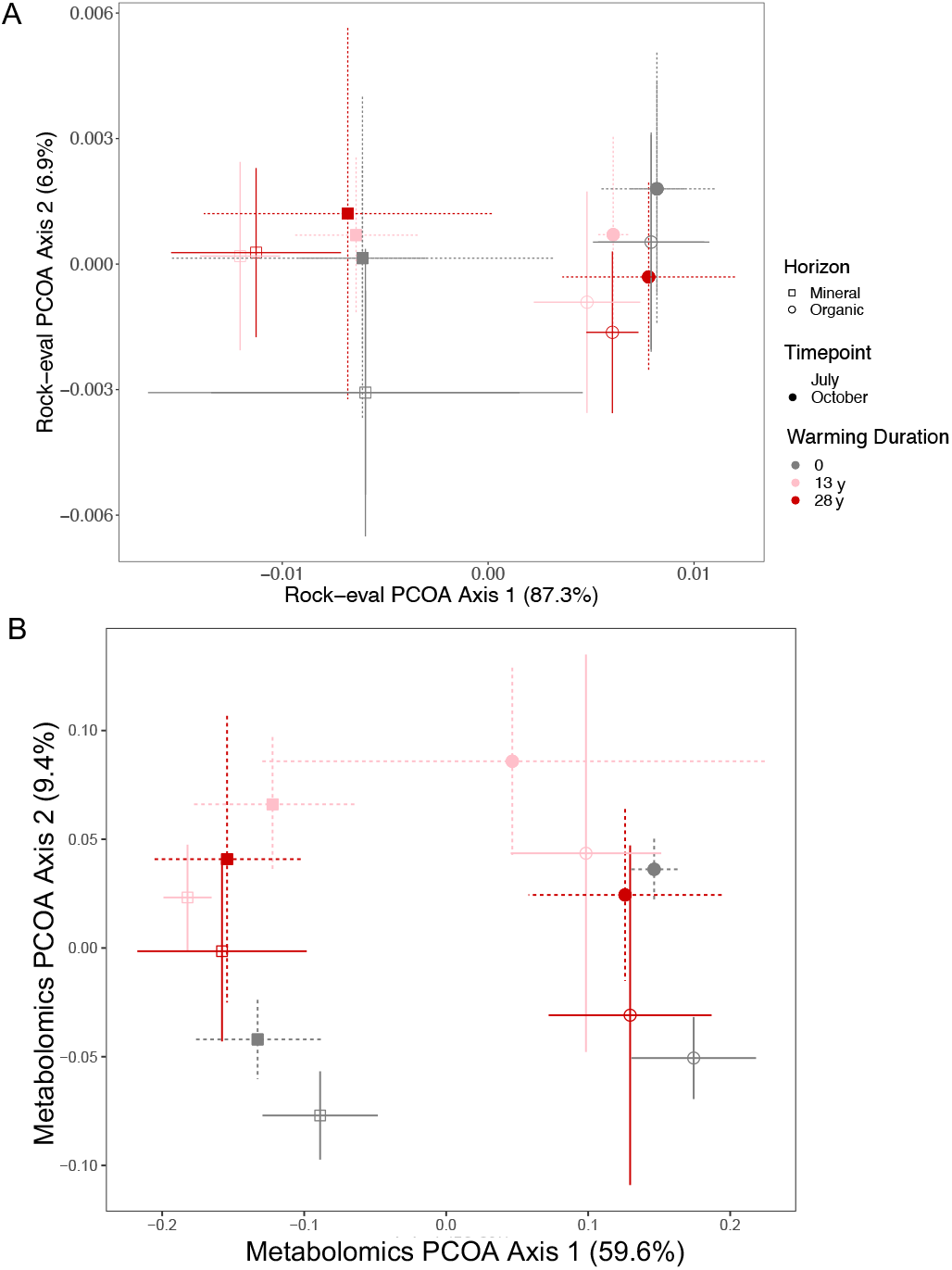
Soil organic matter quality. Ordination of soil organic matter composition from control and heated soils differing in warming duration (13 or 28 years) sampled in July and October based on Principal COordinate Analysis (PCOA) from the pyrolyzed fraction of SOM based on Rock-Eval^®^ (A) and water-extractable polar metabolites (B).

The large variation in the ^18^O signal detected in DNA meant that growth and CUE data did not fit the expected unimodal or monotonic decline pattern, and we were unable to fit quadratic curves to the data to assess subtle shifts in the temperature optimum of growth or CUE. However, median CUE was greatest at 15°C across July samples and 15 or 20°C in October samples (Figure 3).

### Microbial activity reflects soil organic matter chemistry

To evaluate changes in microbial physiology we used correlations between microbial activity per unit microbial biomass and soil properties. This allowed us to evaluate how processes rate and temperature sensitivity of microbial activity is associated with SOM chemistry (Figure 5a). The temperature sensitivity of microbial activity was generally positively correlated with the concentration of total and dissolved soil organic matter, and the thermal lability of soil organic matter, and negatively correlated with the NOSC and chemical complexity of WEOM (Figure 5a). Microbial biomass-specific respiration rate at 20°C showed similar correlations to SOM indices as the temperature sensitivity, but the rates of extracellular enzyme activity showed contrasting patterns. Notably, mass-specific oxidative enzyme activity was greatest in soils characterized by chemically-complex and high NOSC WEOM and with low SOM lability indices (Figure 5a). We also observed a significant positive correlation between the temperature sensitivity of respiration and the temperature sensitivity of BG enzyme activity (Figure 5b), indicating that soils with a pool of cellulolytic enzymes capable of responding to increasing temperatures also respired more in response to changing temperatures. The same was not true for the amino sugar degrading enzyme NAG, whose temperature sensitivity was uncorrelated with that of respiration (Supplementary Figure S6).

**Figure 5.**
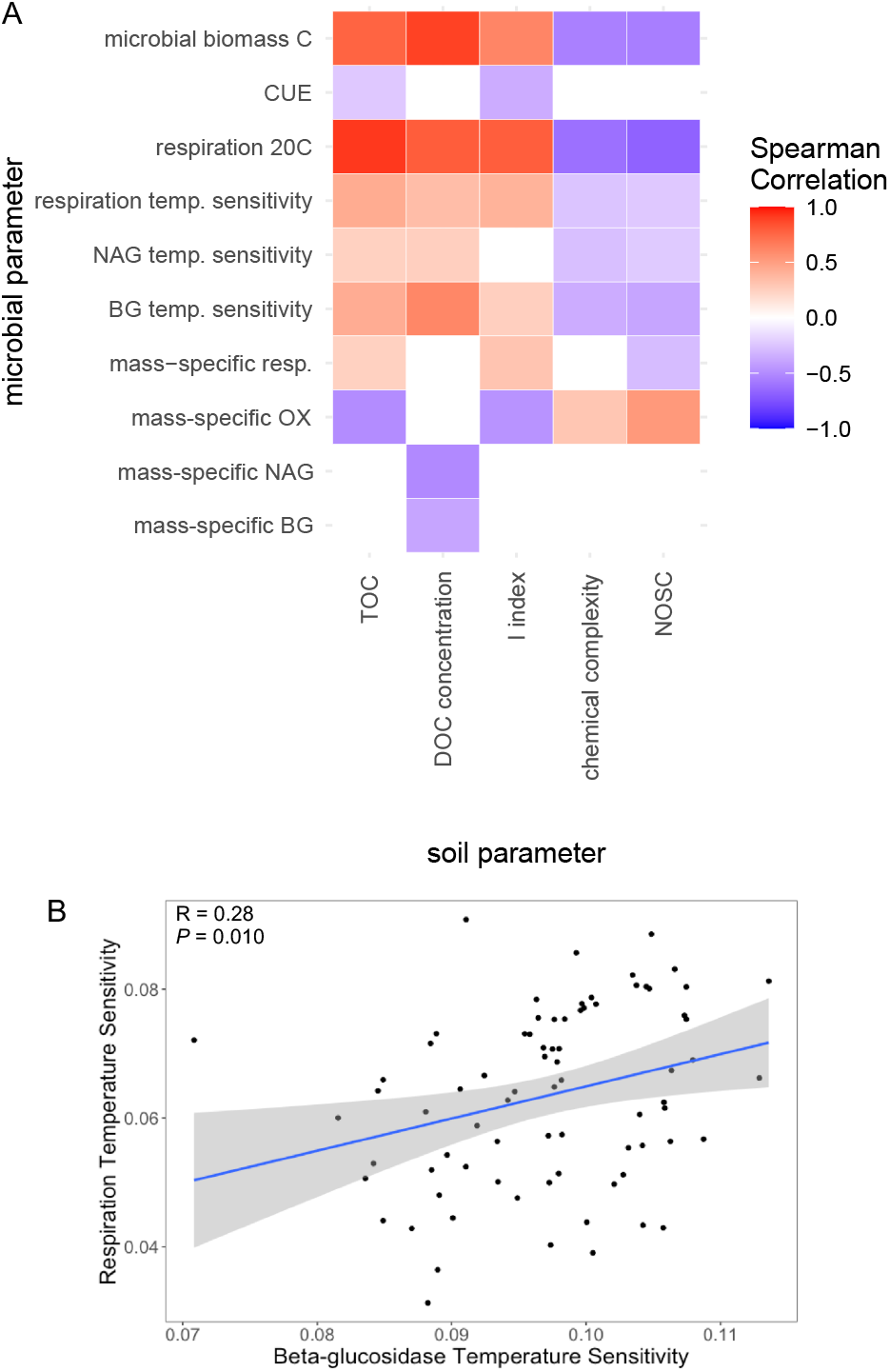
Relationship between the quality of soil organic matter and the response of microbial physiology. Heat map of Spearman correlation coefficients between soil parameters and microbial physiology measurements across all samples (A), and relationship between the temperature sensitivity of respiration and extracellular enzymatic activity (B).

### Effects of warming on microbial physiology play out through its effects on soil C

We used path analysis to integrate the relationships between the abiotic and biotic measurements performed here and test the hypothesis that long-term warming drives microbial activity through its effect on soil C. The path analysis confirms this hypothesis and shows that long-term warming treatment, season and soil horizons influenced the carbon quantity and quality in the soil while also affecting respiration temperature sensitivity (Figure 6). However, long-term warming duration has no direct effect on microbial physiology or WEOM and SOM quality. Unexpectedly, the nominal oxidation state of water extractable organic matter was not a key integrator of shifts in SOM chemistry and did not drive respiration temperature sensitivity (Figure 6). Rather, temperature sensitivity of respiration was directly correlated with quantity of carbon and the size and temperature sensitivity of the extracellular enzyme pool, and indirectly with SOM quality measured by Rock-Eval^*®*^ (Figure 6). Season had a direct effect on the dissolved fraction of SOM and C quality while increasing soil depth decreased the SOM quality (e.g. labile fraction). Respiration temperature sensitivity was driven directly by the enzyme pool in soils, carbon quantity and the enzyme temperature sensitivity. The soil potential extracelullar enzyme activity was driven by C quantity and season while the enzyme temperature sensitivity was driven by both season and carbon quality. Although microbial community can modulate thermal acclimation, the fungal to bacterial ratio was only driven by soil horizon and did not influence other parameters in our model. Our SEM model explained 28% of the variance in the temperature sensitivity of respiration, though many of the drivers of temperature sensitivity were explained to a much greater degree.

**Figure 6.**
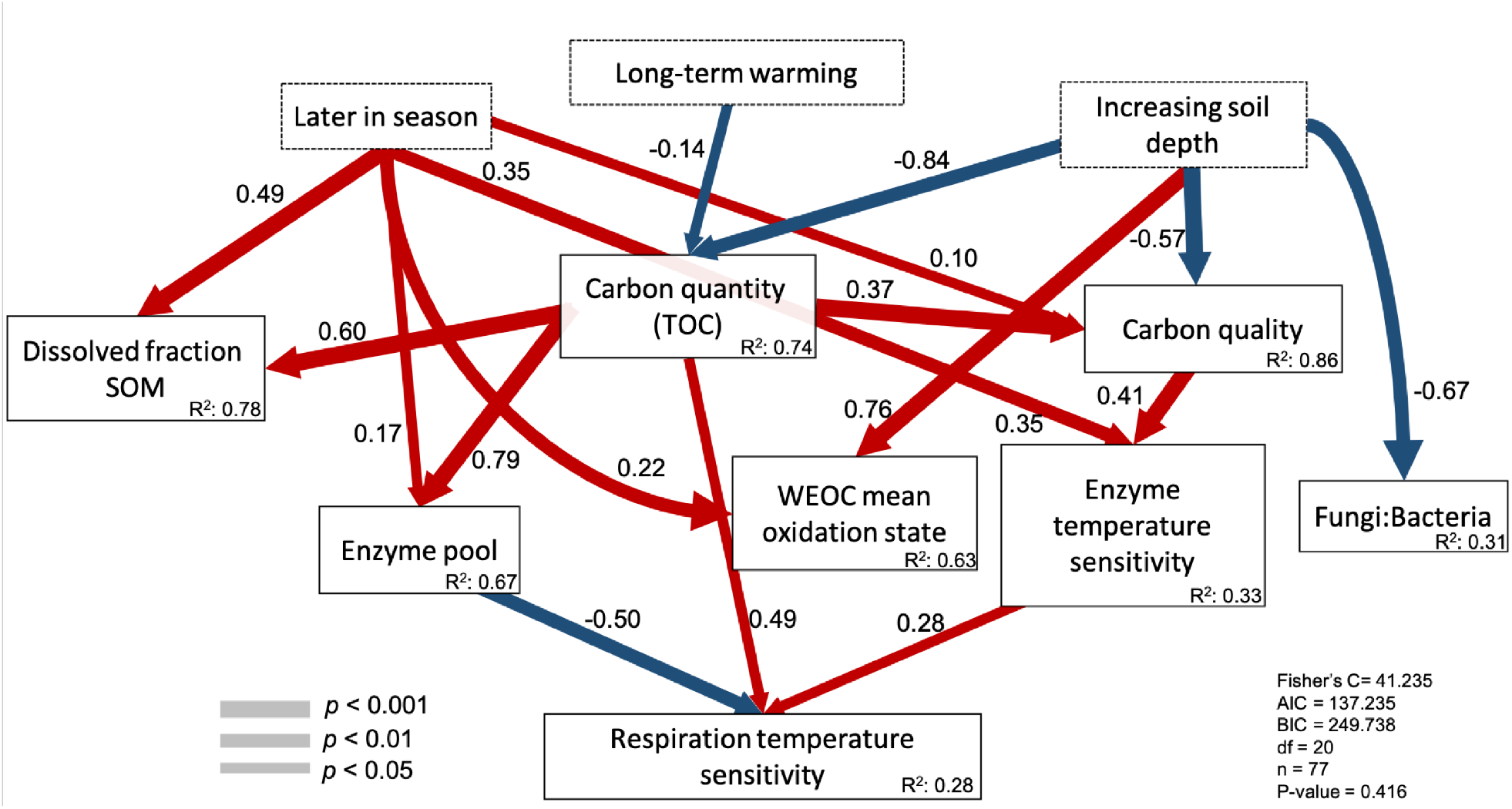
Structural equation model showing the relative influence of long-term warming duration, soil horizons and different seasons (July and October) on abiotic factors and components of microbial physiology driving the respiration temperature sensitivity. Significant paths are shown in red if positive or in blue if negative with their path coefficients. Path width corresponds to degree of significance as shown in the lower left. The amount of variance explained by the model (R^2^) is shown for each response variable, and measures of overall model fit are shown in the lower right. Dissolved fraction of SOM: DOC measured by a TOC-L analyzer; carbon quantity: total carbon quantified during rock-eval; C quality: PCOA axis 1 of rock-eval thermal pyrolysis; enzyme pool: composite variable of maximum activity recorded for BG, NAG and oxidative enzymes at 20°C; WEOM mean oxidation state: SOM oxidation state calculated from polar metabolites; enzyme temperature sensitivity: composite variable of BG, NAG and oxidative enzymes maximum activity response to temperature while incubated from 4°C to 30°C; Fungi:Bacteria: 16S rRNA gene copy number g^−1^ soil: ITS gene copy number g^−1^ soil; respiration temperature sensitivity: respiration response to temperature while incubated from 4°C to 30°C. Global goodness-of-fit: Fisher’s C.

Path analysis of CUE at 20°C shows that it is directly affected by respiration and growth as expected, but CUE is also driven by other factors such as the dissolved SOM fraction and season, with higher CUE measured in October compared to July (Figure 7). The mass-specific microbial respiration component of CUE was much better explained than the growth component R^2^=0.77 vs. 0.14), with 86% of the variation in CUE explained overall (Figure 7). This was expected given the large variation in our measurements of growth rate. Distinct factors also drove respiration and growth differently. The mass-specific enzyme pool only drives respiration while carbon quality correlates positively with mass-specific respiration and negatively with mass-specific growth.

**Figure 7.**
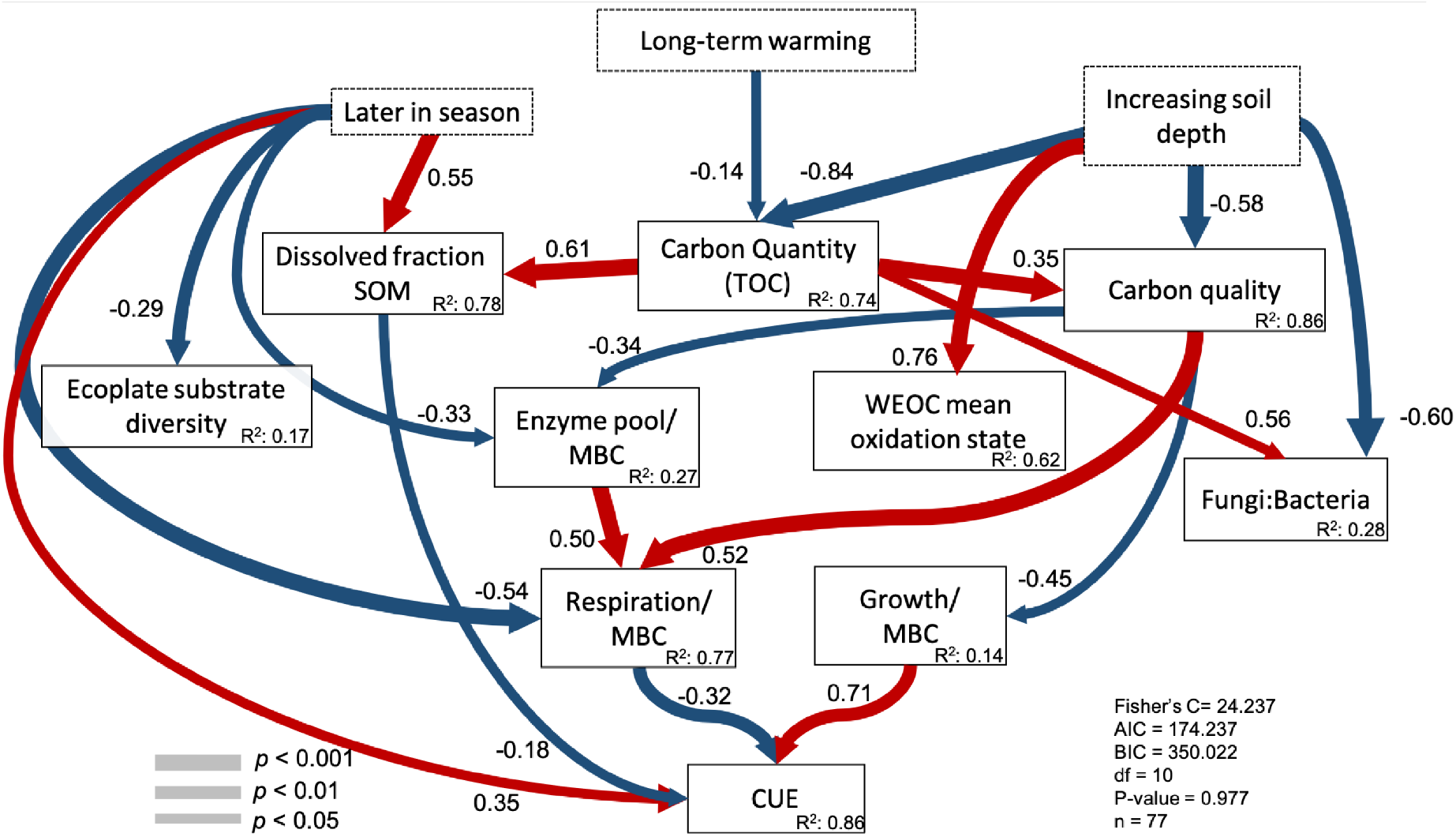
Structural equation model showing the relative influence of long-term warming duration, soil horizons and different seasons (July and October) on abiotic factors and components of microbial physiology driving CUE. Significant paths are shown in red if positive or in blue if negative with their path coefficients. Path width corresponds to degree of significance as shown in the lower left. The amount of variance explained by the model (R^2^) is shown for each response variable, and measures of overall model fit are shown in the lower right. Dissolved fraction of SOM: DOC measured by a TOC-L analyzer; carbon quantity: total carbon quantified during rock-eval; carbon quality: PCOA axis 1 of rock-eval thermal pyrolysis; enzyme pool/MBC: composite variable of maximum activity recorded for BG, NAG and oxidative enzymes by microbial biomass carbon; SOM mean oxidation state: WEOM oxidation state calculated from polar metabolites; Fungi:Bacteria: 16S rRNA gene copy number g^−1^ soil: ITS gene copy number g^−1^ soil; respiration/MBC: respiration measured at 20°C/ microbial biomass carbon; growth/MBC: growth measured at 20°C/ microbial biomass carbon and CUE: carbon use efficiency measured at 20°C. Global goodness-of-fit: Fisher’s C.

In order to avoid the potential problems associated with predicting CUE from its constituent components (i.e. growth and respiration), we also built a SEM without these variables to evaluate how much the other variables we measured directly explained CUE (Supplementary Fig S12). This model captures 39% of CUE variance and the direct drivers were season, mass specific enzyme pool and carbon quality. Interestingly, high quality carbon is associated with lower CUE in both models because it accelerates mass-specific respiration while slowing mass-specific growth (Figure 7 and Supplementary Figure S12).

## Discussion

Here, we evaluated the hypothesis that the effect of warming on microbial physiology plays out through its effects on the quantity and quality of soil organic matter. We found that while soil chemistry had predictable correlations with microbial activity, the effects of warming on soil organic matter chemistry — and, in turn, microbial physiology — were limited to summer. Therefore, the effect of warming on microbial physiology depends on its effects on SOM chemistry.

The effect of warming on SOM quality was most apparent in the mineral soil, where soil lability index was reduced (Figure 2b). This is consistent with previous studies at these sites which show that warming-associated changes in soil organic matter quality are especially apparent in the mineral soil, where typical plant-derived compounds such as lignin and polysaccharides appeared more depleted after 4 and 23 years of warming (Pisani, Frey, Simpson, & Simpson, 2015; Pold et al., 2017). Greater decomposition has increased overall relative abundance of lipids (Pec et al., 2021; Pold et al., 2017; Pisani et al., 2015; VandenEnden et al., 2021) despite a reduction in plant lipids (Pisani et al., 2015) under warming at these sites, indicating further microbial processing. Lignin and lipids are both among the least oxidized (LaRowe & Van Cappellen, 2011) and most readily-pyrolyzed (Sanderman & Grandy, 2020) compounds found in soil, such that a potential shift towards being more lipid-rich and lignin-poor under warming in the organic soil would not be detected. However, it is more probable that any signal of increased microbial processing was drowned out in the high plant input:microbial processing ratio in the organic soil. This is consistent with the observation that warming does not consistently alter SOM quality throughout seasons, even in the more microbial necromass-dominated mineral soil (Fig 2b and Supplementary Fig. 1a); differences were only apparent in our summer sampling when belowground inputs likely dominated carbon inputs (Abramoff & Finzi, 2015; Cheeke et al., 2021), and not following the autumn leaf litter influx. This is consistent with a wealth of studies indicating that season strongly regulates the input of plant material and nutrients in temperate forest soils and that this seasonality is associated with variation in microbial activity and biomass (Cheeke et al., 2021; Kaiser et al., 2010; Pold et al., 2017; Rasche et al., 2011; Wardle et al., 2004).

This seasonal dependence of warming effects on SOM chemistry was further associated with temporally-inconsistent effects of warming on microbial physiology. Limited substrate availability has been cited as necessary condition for microbial community physiological acclimation under warming (Bradford, 2013; Kirschbaum, 2004; Alster et al., 2022; Luo, Wan, Hui, & Wallace, 2001; Fierer, Colman, Schimel, & Jackson, 2006; Moinet et al., 2021). We only observed apparent thermal acclimation of C cycling processes when the input of fresh plant material was limited (i.e. in summer), and not once a pulse of fresh aboveground plant input reaches the soil (i.e. in autumn; Figure 2b and 3). Temperature is an important driver controlling how soil microbes degrade organic matter (Bradford, 2013), and to a certain degree elevated temperatures may help microbes overcome substrate limitation by accelerating the rate at which extracellular enzymes process SOM. However, as warmer temperatures sustain accelerated microbial activity through time, ultimately substrates become depleted, explaining the often-observed only temporary increase in soil respiration under chronic warming (Luo et al., 2001; Walker et al., 2018; Romero-Olivares, Allison, & Treseder, 2017). This substrate depletion leads to a reduction in microbial biomass followed by a decrease in microbial activity (Walker et al., 2018). However, as seen here, seasonal flushes of fresh aboveground litter inputs may temporarily mitigate this warming-induced substrate depletion, allowing microbial biomass and therefore process rates to somewhat recover to the levels observed in the absence of chronic warming. This concept of substrate limitation induced reductions in microbial activity via reductions in microbial biomass is supported by the observation that mass-specific C processes rates are less affected by warming compared to the activities measured on a per gram of soil basis (Figure 2b). Accordingly, two metrics strongly correlated with microbial biomass - total organic carbon and size of the extracellular enzymatic pool - along with temperature sensitivity of extracellular enzyme activity, were the main drivers of respiration temperature sensitivity.

Part of the greater respiration temperature sensitivity with larger enzyme pools may be linked to greater temperature sensitivity of extracellular enzyme activity, enabling greater supply rate of products for microbial metabolism without greater investment in those enzymes. However, the connection between the temperature sensitivity of respiration and extracellular enzyme activity is not equally linked to all SOM pools equally; soils with a bigger pool of cellulolytic enzymes are more capable of responding to increasing temperatures and consequently respired more in response to changing temperatures, but the same was not true for soils with high NAG activity. Therefore, our results implicate the interaction between the quantity and quality of SOM in driving microbial community activity (Kirschbaum, 2004; Moinet et al., 2021) and functioning (Domeignoz-Horta et al., 2021).

Carbon use efficiency is an important integrator of C quality with microbial metabolism, both because of the costs associated with extracellular enzyme production necessary to access those substrates (Domeignoz-Horta et al., 2020; Malik, Puissant, Goodall, Allison, & Griffiths, 2019), and because of differences in the communities able to use and the energetic yield associated with the use of those different substrates (Pold, Domeignoz-Horta, Morrison, et al., 2020). The respiration component of carbon use efficiency was more responsive to changes in SOM and WEOM pools compared to growth, indicating that growth has a relatively fixed cost compared to respiration. This is in line with the observation that microbes may downregulate the protein production machinery in response to warming-induced substrate depletion, allowing them to sustain growth (Söllinger et al., 2022) while the respiration component could then absorb changes in substrate quantity and quality. This downregulation of the protein machinery does not require a change in the active community (Söllinger et al., 2022). Changes in the microbial community have historically been variable and limited at our study sites (Roy Chowdhury et al., 2021; DeAngelis et al., 2015; S. Frey, Drijber, Smith, & Melillo, 2008; Pec et al., 2021), and we did not observe a change in the F:B ratio under warming here. Nor did we observe a correlation between F:B ratio and CUE. This reinforces the idea that substrates rather than microbial identity have the dominant effect on CUE and its response to long-term warming in these soils.

Microbial activity measurements are more strongly correlated with the thermal-lability (I index) of the total organic matter measured by Rock-Eval^*®*^ than the water-extractable pool measured by polar metabolomics. Regardless of whether this is due to the latter method missing the compounds most important for microbial activity over the course of the assays period, or whether the chemistry of bulk SOM rather than the water-extractable component is truly a better predictor of microbial physiology, our results show that SOM thermal-lability index based on Rock-Eval^*®*^ captures organic matter pools which vary between soils characterized by divergent microbial community metabolisms. Nonetheless, the warming impact on resource availability is inconsistent due to seasonality in inputs (Figure 2 and 3), which might result in a variable selection regime (McDonald, 2019) limiting the degree of intrinsic thermal acclimation of the microbial community to long-term warming in these soils. As such, our results are consistent with previous findings that the response of microbial communities to global warming and its potential impact on soil carbon stocks will depend on concurrent changes in soil organic matter chemistry and availability to microorganisms (Bradford et al., 2008; J. M. Melillo et al., 2017). While we still lack a clear understanding of the controls that drive variation in the response of soil carbon to warmer temperatures globally (Van Gestel et al., 2018), our results identify the direct drivers of apparent thermal acclimation of C processes drawing attention to the importance to monitor soil C chemistry to better understand possible soil-carbon feedback mechanisms to the atmosphere as the world warms.

## Conclusion

Here, we evaluated the interplay between the biotic and abiotic components of an ecosystem exposed to long-term warming to understand the mechanisms underlying the microbial physiological response to warming. We recorded that the apparent thermal acclimation of respiration and extra-cellular enzyme activities in heated soils compared to control soils in the summer sampling was caused by lower substrate availability due to long-term warming. We did not detect temperature sensitivity in any C-cycling process in autumn, likely due to litter deposition alleviating summer substrate depletion. By disentangling the direct drivers of apparent thermal acclimation in long-term warmed soils these results highlight that if we are to improve soil C predictions in a warming planet we need to account for concurrent changes in soil carbon quantity and quality.

## Acknowledgements

Funding for this project was provided by the National Science Foundation (NSF)(DEB-1749206) to K.M.D.; NSF Long-Term Research in Environmental Biology (DEB-1456528) to K.M.D, S.D.F. and J.M.M.; and NSF Long-Term Ecological Research (DEB-1832210) programs. This work was also conducted with support (proposal 506489: 10.46936/10.25585/60001340 to L.A.D-H and K.M.D.) from the U.S. Department of Energy Joint Genome Institute (https://ror.org/04xm1d337), a DOE Office of Science User Facility, is supported by the Office of Science of the U.S. Department of Energy operated under Contract No. DE-AC02-05CH11231.

## Author contributions statement

G.P., L.A.D-H. and K.M.D. designed lab experiments. S.D.F and J.M.M. designed field experiments. G.P. L.A.D-H., M.K. and H.E. conducted the experiments. L.A.D-H., G.P., D.S, K.L. analyzed data. G.P. and L.A.D-H. wrote the first draft of the paper. G.P., L.A.D-H, K.M.D, S.D.F, J.M.M., D.S., K.L., E.V., H.E., T.N., E.E-F., C.P. and M.A.K contributed to revise the manuscript.

## Additional information

### Accession codes

Metagenomes generated during this study are available at the JGI Gold portal with the ID Gs0149985 (Supplementary table 3). All data and codes used in this study can be obtained in the Open Science Framework Project (https://osf.io/zdhfb/) and from the authors upon request.

