## Supplementary Material for "Substrate availability and not thermal-acclimation controls microbial temperature sensitivity response to long term warming"

### Supplementary Figures

**Supplementary Table 1:** Basic site characteristics of experiments used in this study.

| Characteristic | SWaN plots | Prospect Hill |
|---|---|---|
| **Latitude, longitude** | 42.54°N, 72.18°W | 42.54°N, 72.18°W |
| **Year started** | 2006 (13 years) | 1991 (28 years) |
| **Years of warming at time of sampling** | 13 years | 28 years |
| **Plot size** | 3 * 3m | 6 * 6m |
| **No. of plots sampled** | 5 | 5 |
| **Soil pH, O-horizon** | 3.72 | 3.82 |
| **Soil pH, 0- to 10-cm mineral** | 4.38 | 4.41 |
| **mean O horizon depth sampled +/- SE: range (cm)** | 5.6 (1.6); 1.5-11.2 | 3.5 (0.4); 2.4-5.5 |
| **Total soil C (mean +/- SE) (g m-2)** | | |
| **O-horizon** | 3,314 (404) | 2,565 (247) |
| **0- to 10-cm mineral** | 3,478 (121) | 2,859 (444) |
| **Soil series** | Gloucester | Gloucester |
| **Dominant overstory vegetation** | *Acer rubrum, Acer pensylvanicum, Betula papyrifera, Fagus grandifolia, Quercus rubra, Quercus velutina* | *A. rubrum, A. pensylvanicum, B. papyrifera, Q. velutina* |

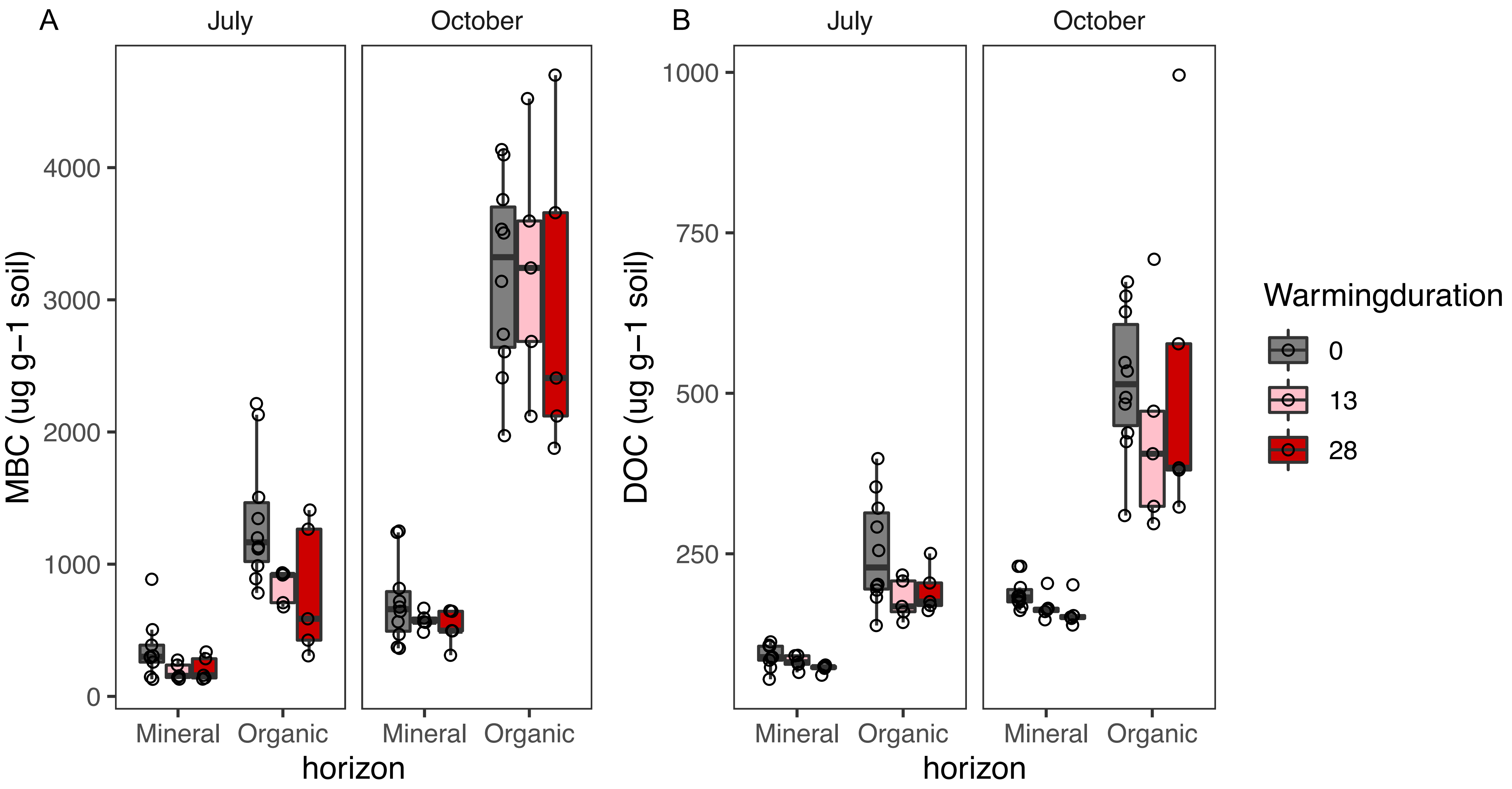


**Supplementary Figure 1.** Microbial biomass carbon (A) and dissolved organic carbon (B) response to seasons, soil depth and long-term warming.

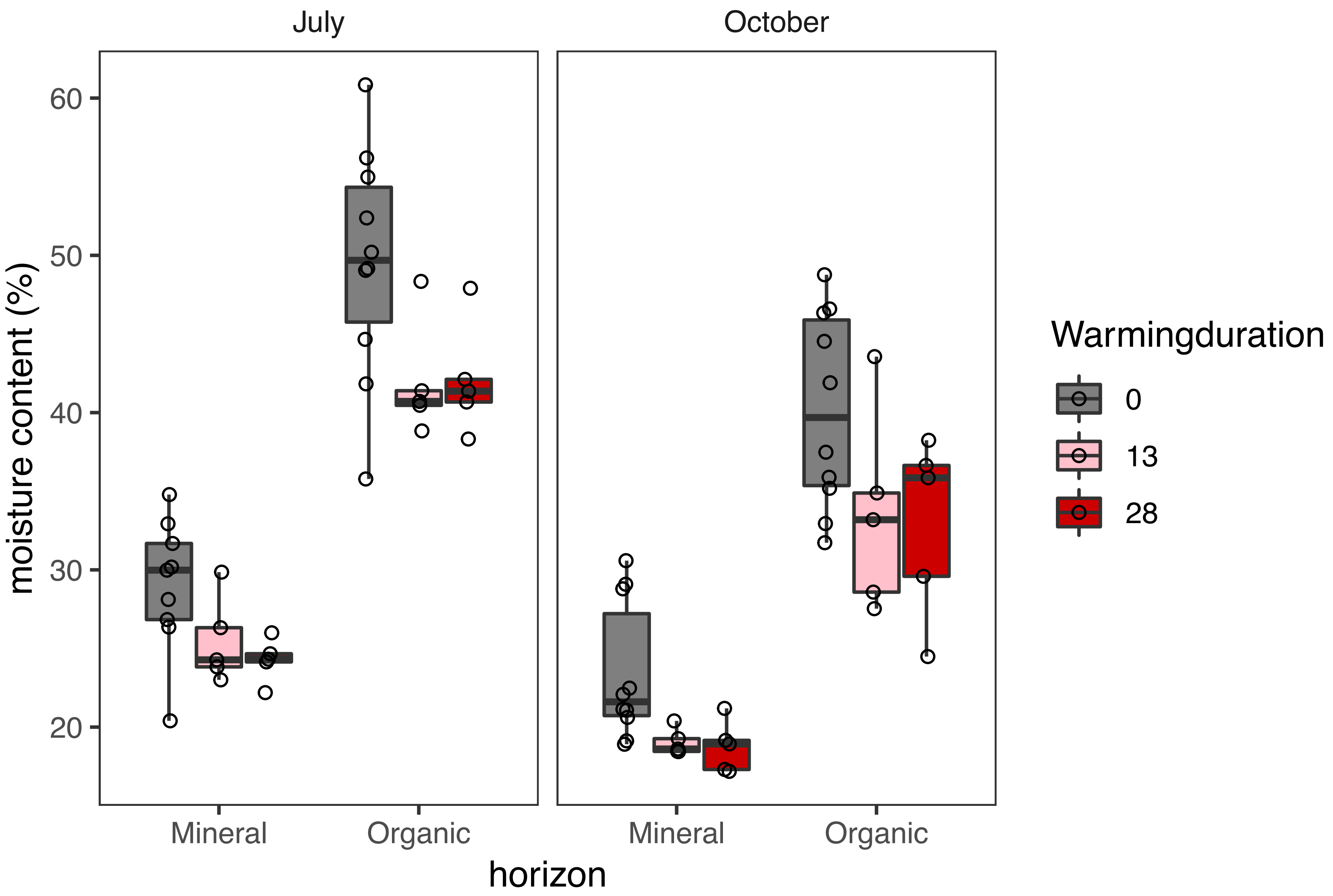


**Supplementary Figure 2.** Soil moisture response to seasons, soil depth and long-term warming.

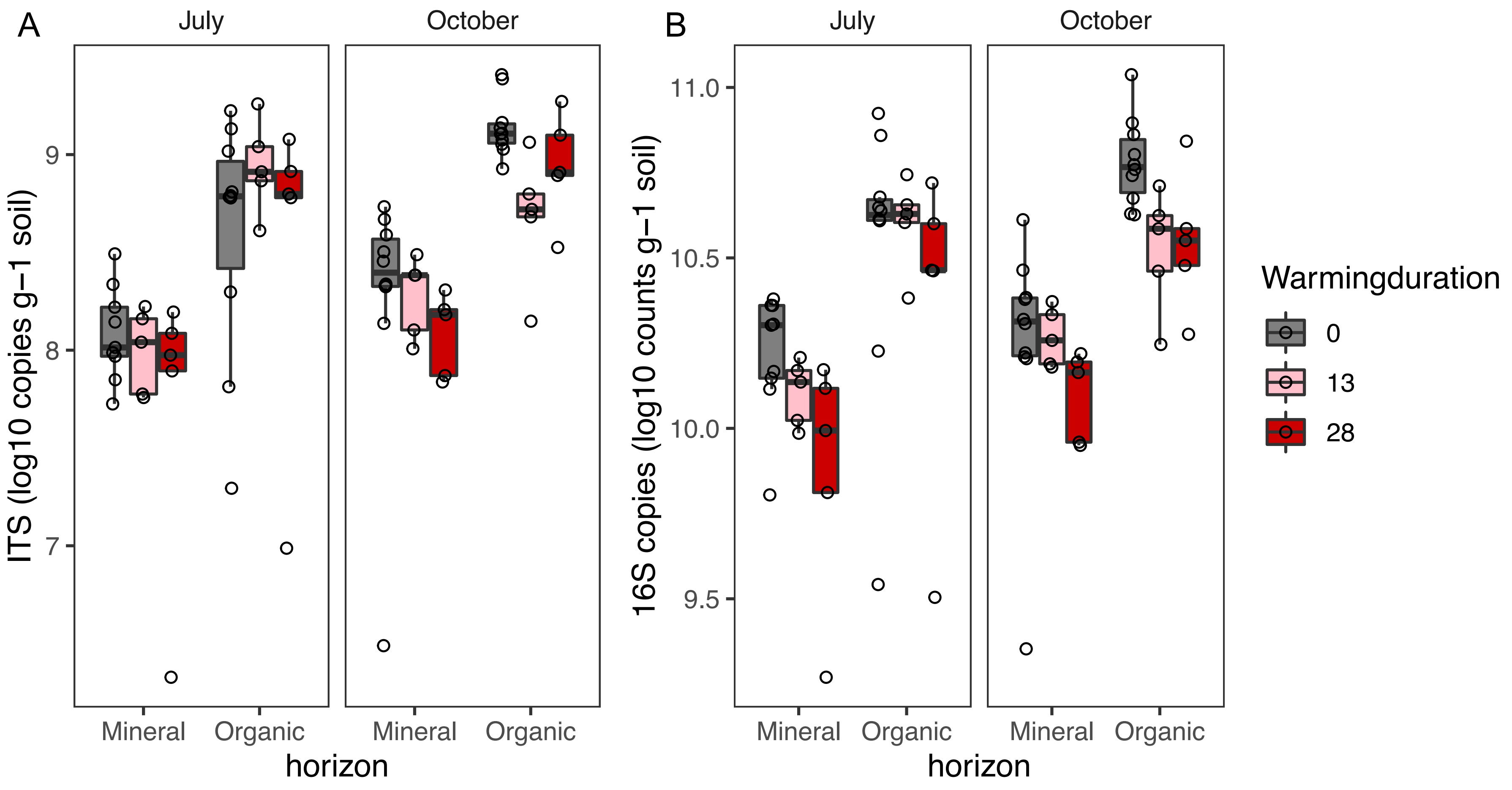


**Supplementary Figure 3.** Fungal (A) and bacterial (B) abundance response to seasons, soil depth and long-term warming.

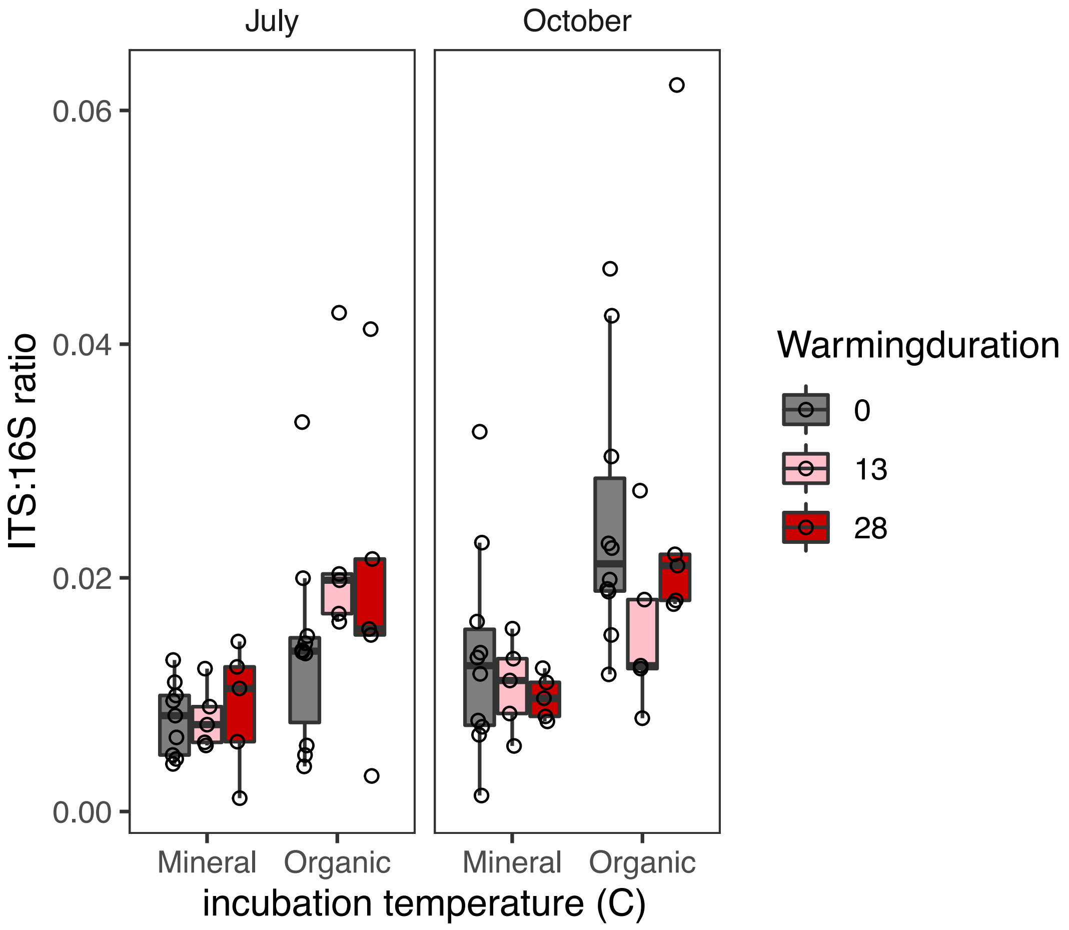


**Supplementary Figure 4.** Fungal to bacterial ratio response to seasons, soil depth and long-term warming.

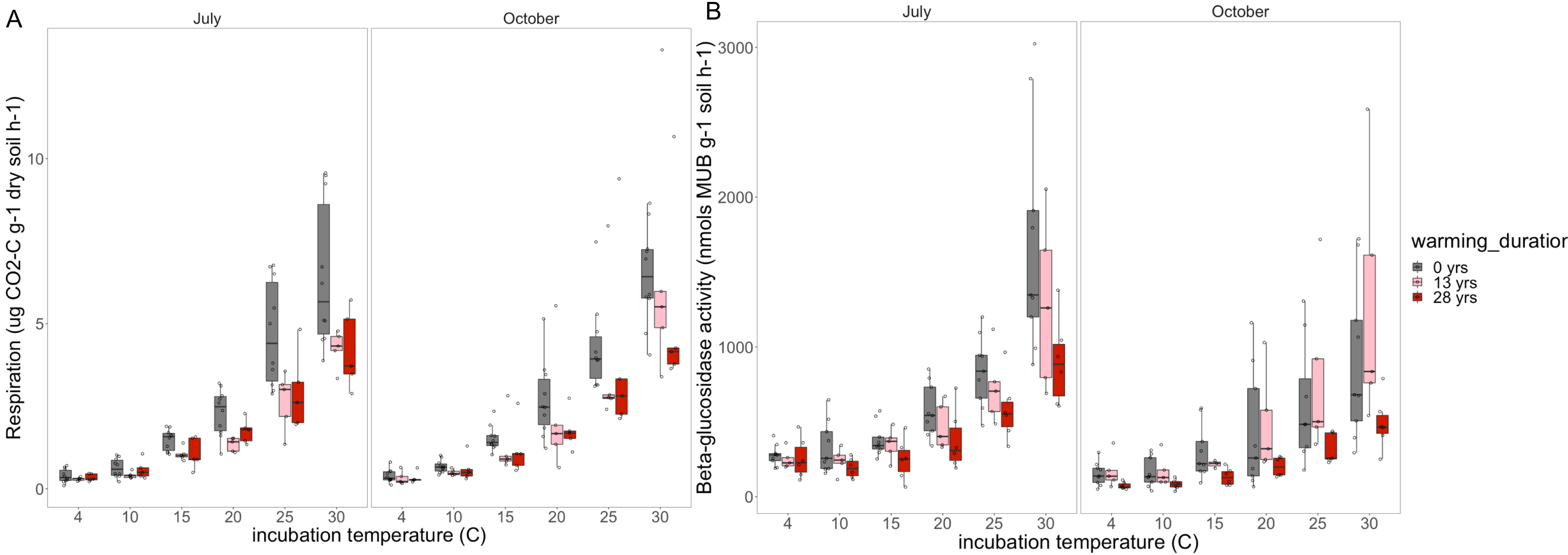


**Supplementary Figure 5.** Temperature sensitivity of respiration (A) and betaglucosidase activity (B) in organic soils measured at different temperatures from 4-30$^{o}$C during laboratory incubations in July and October.

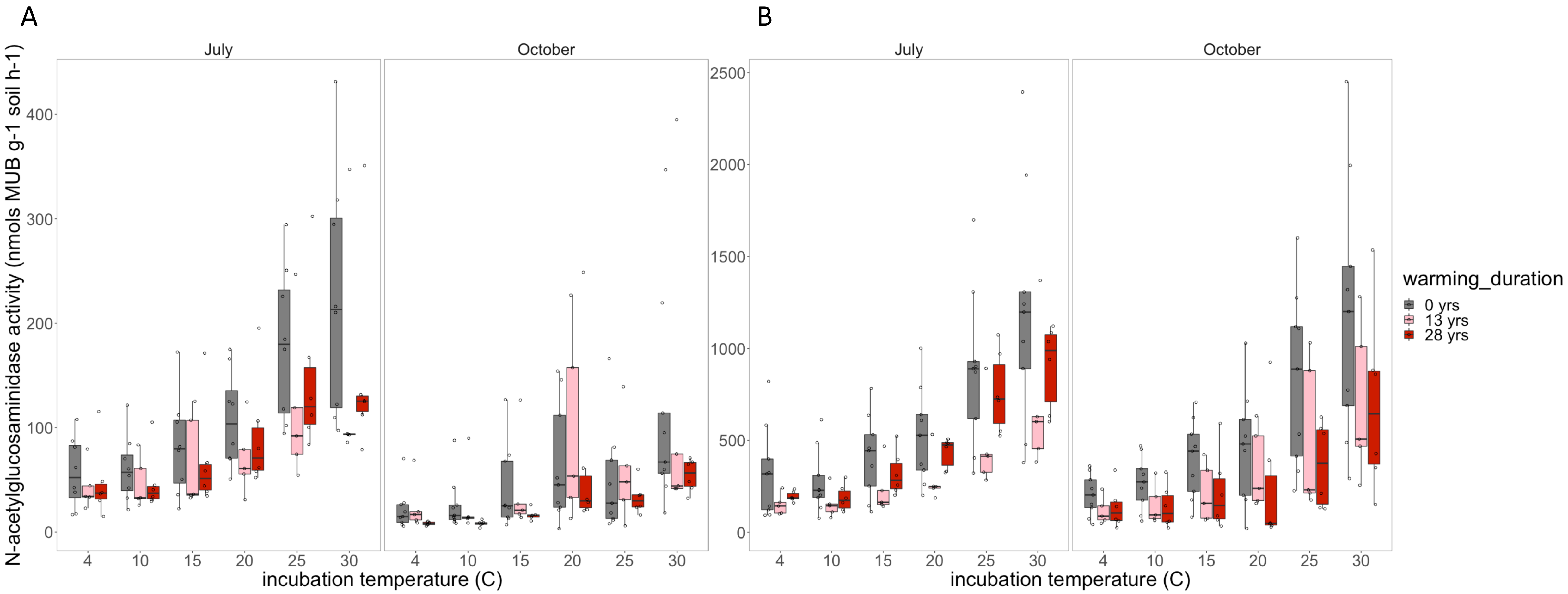


**Supplementary Figure 6.** Temperature sensitivity of N-acetylglucosaminidase in mineral (A) and organic (B) soils. measured at different temperatures from 4-30$^{o}$C during laboratory incubations in July and October.

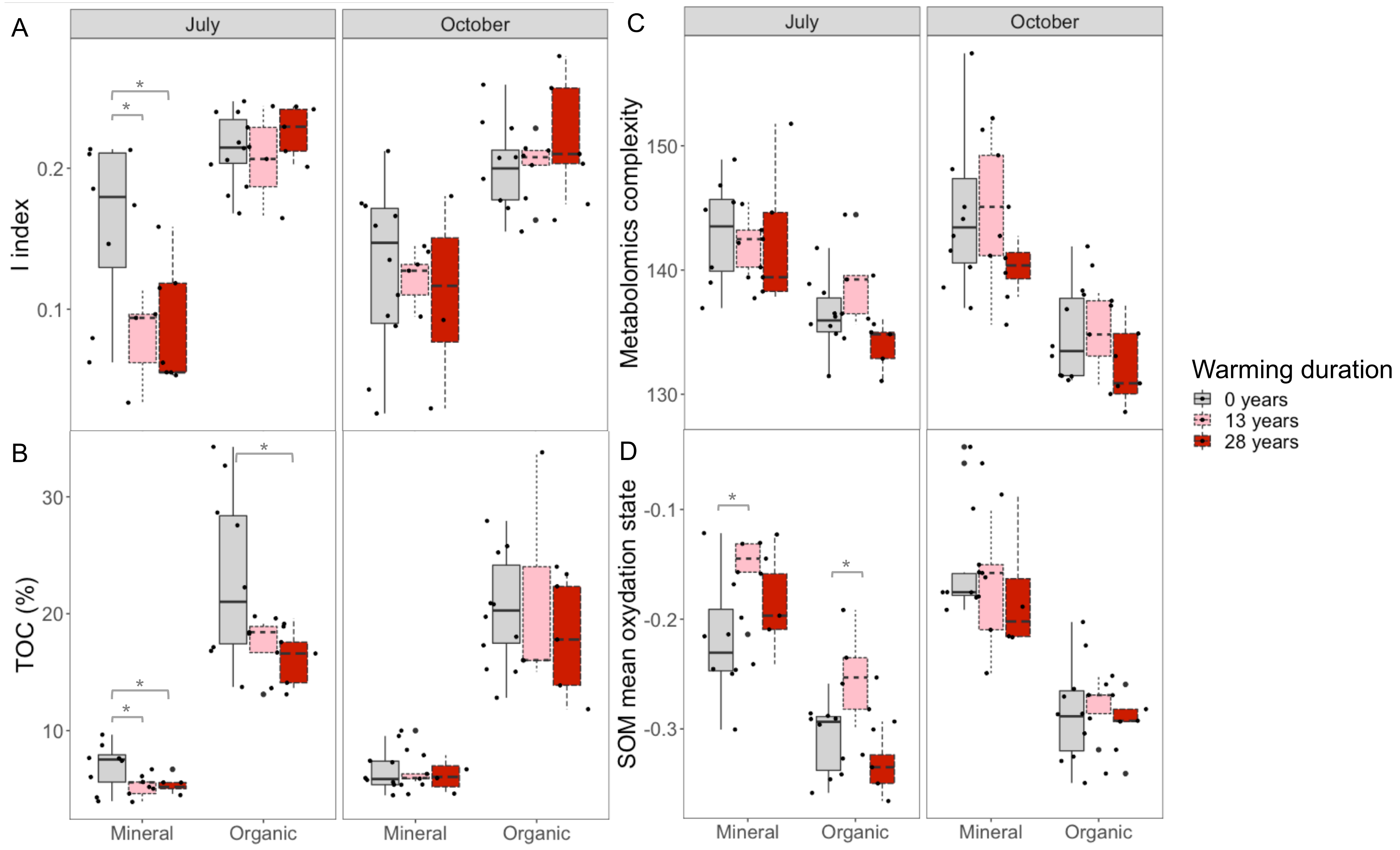


**Supplementary Figure S7:** Soil organic matter quantity and quality. Soil organic matter I-index based on rock-eval (A), total organic carbon (B), Metabolomics complexity (C) and SOM mean oxidation state (D).

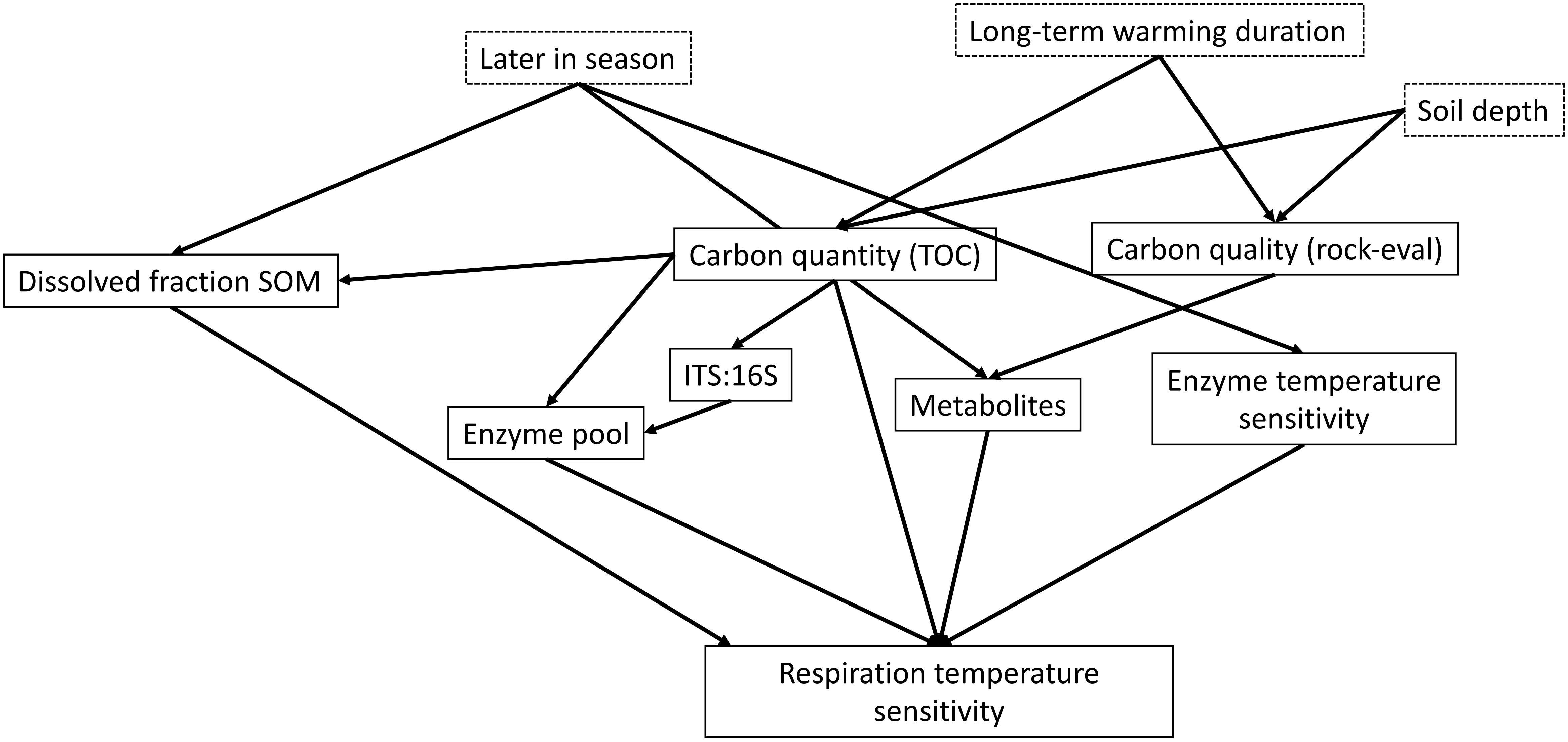


**Supplementary Figure S10.** Hypothesized structural equation model for the respiration temperature sensitivity.

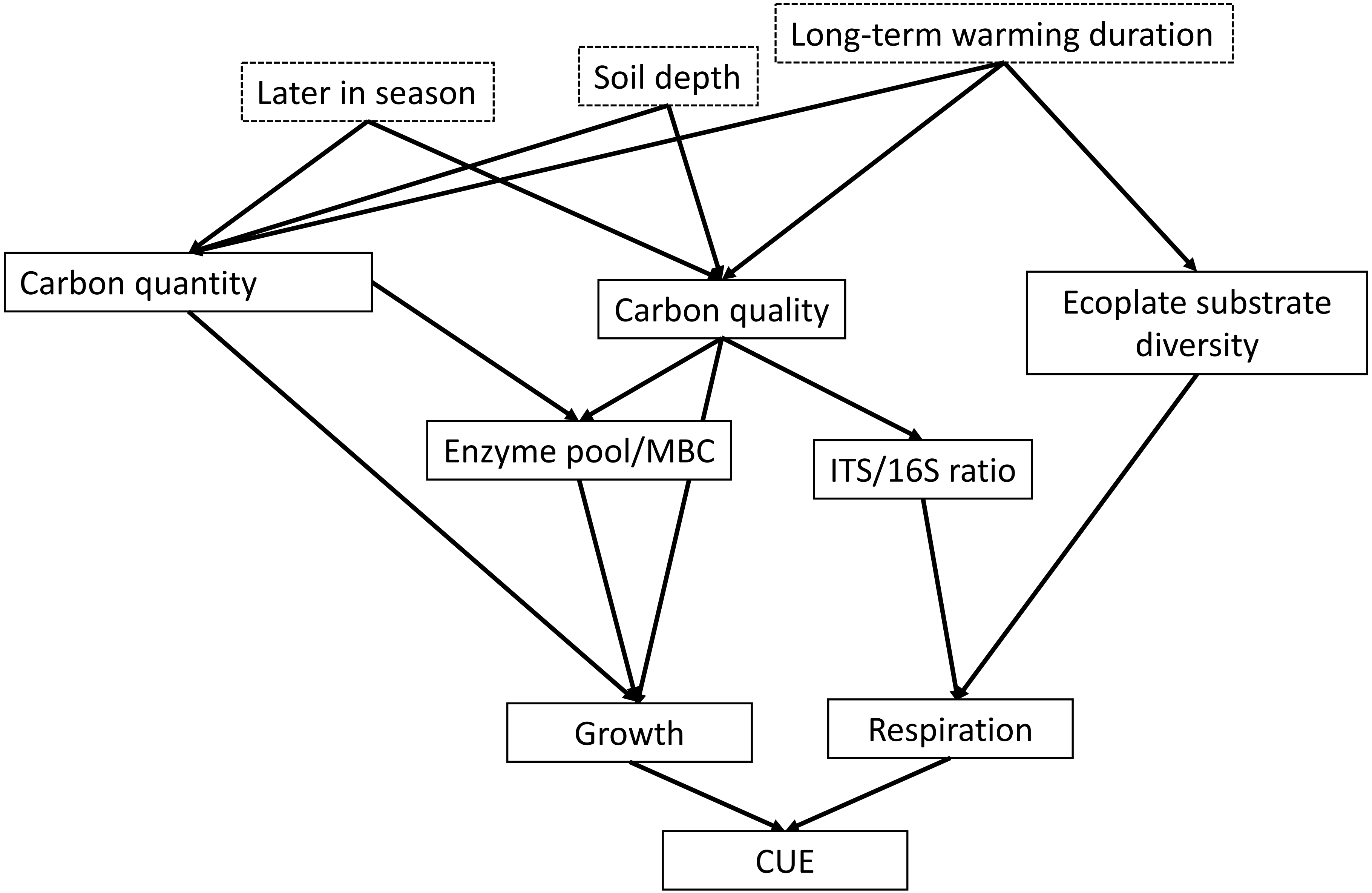


**Supplementary Figure S11.** Hypothesized structural equation model for the CUE.

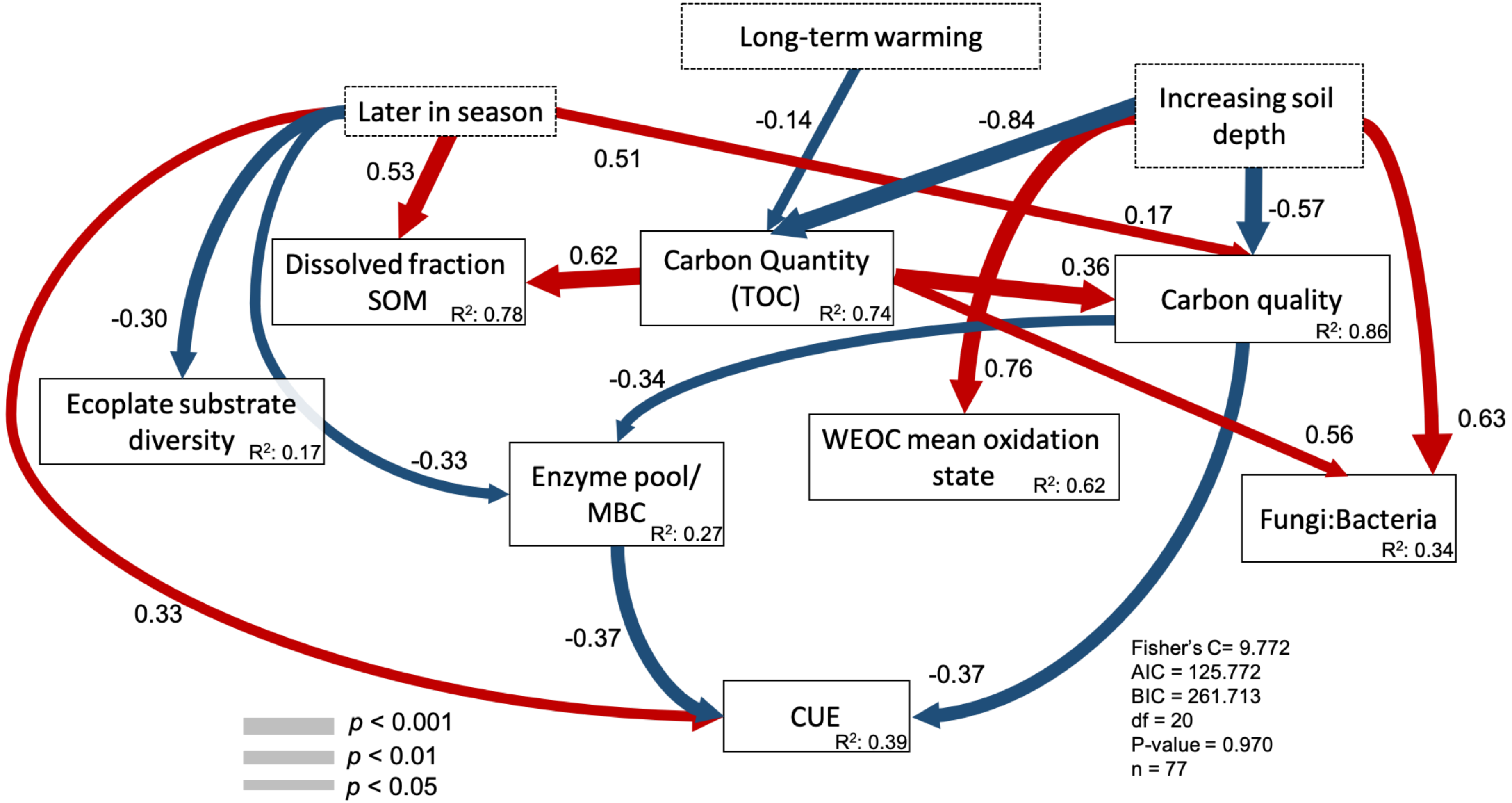


**Supplementary Figure S12. Structural equation model explaining CUE without growth and respiration.** Significant paths are shown in red if positive or in blue if negative. Path width corresponds to degree of significance as shown in the lower left. The amount of variance explained by the model ($R^2$) is shown for each response variable, and measures of overall model fit are shown in the lower right. Dissolved fraction of SOM: dissolved organic carbon measured by a TOC-L analyzer; carbon quantity: total carbon quantified during rock-eval ramped thermal pirolysis; carbon quality: PCOA axis 1 of rock-eval ramped thermal pirolysis; enzyme pool/MBC: composite variable of maximum activity recorded for beta-glucosidase, N-acetylglucosaminidase and oxidative enzymes by microbial biomass carbon; SOM mean oxidation state: SOM oxidation state calculated from polar metabolites; Fungi:Bacteria: 16S rRNA gene copy number g−1 soil: ITS gene copy number g−1 soil; respiration/MBC: respiration measured at 20C/ microbial biomass carbon; growth/MBC: growth measured at 20C/ microbial biomass carbon and CUE: carbon use efficiency measured at 20C. Global goodness-of-fit: Fisher's C.

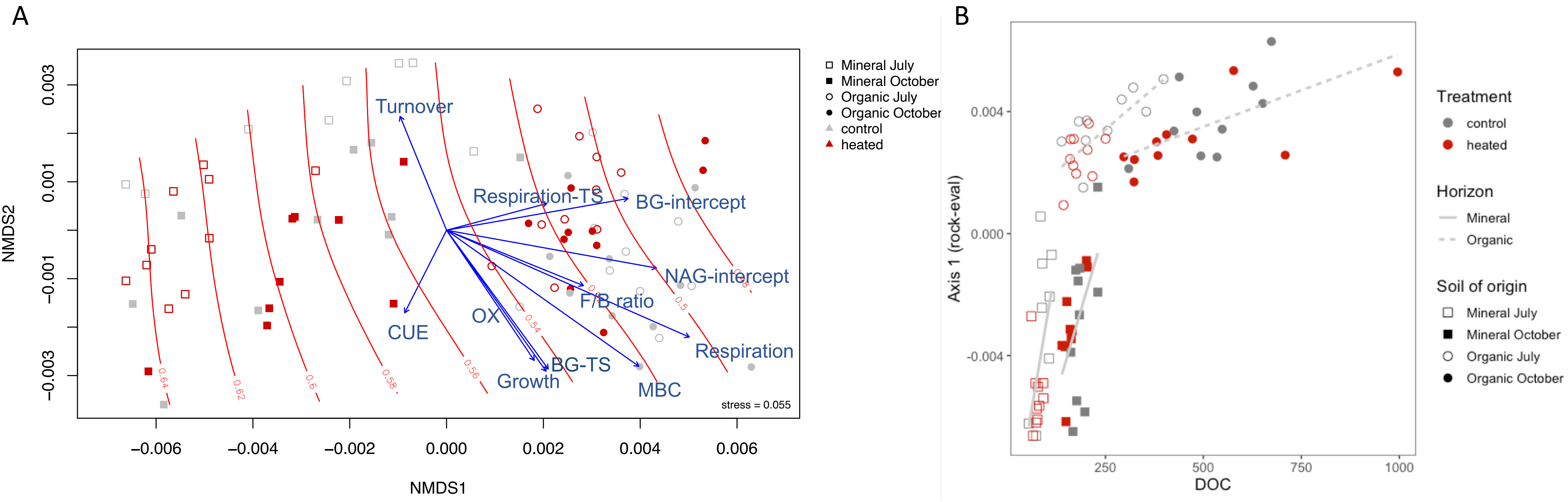


**Supplementary Figure S13. Ordination of soil organic matter composition and relationship between first axis of SOM ordination composition and DOC.** Non-metric multidimensional scaling of Bray–Curtis distance from the pyrolyzed fraction of SOM based on Rock-Eval® analysis. Red contour lines represent the SOM thermal-stability R-index with higher numbers indicating more thermal-stable SOM. Significant explanatory variables ($P < 0.05$) are represented as blue vectors and the lengths of the arrows are proportional to the strength of the correlation; CUE represents the carbon use efficiency; MBC corresponds to microbial biomass carbon (µg C $g^{-1}$ dw soil); turnover the turnover of the community in days; Respiration-TS and BG-TS represents the respiration and beta-glucosidase temperature sensitivity measured during microcosms incubation from 4 to 30$^{O}$C. OX represent maximum oxidative enzymatic activity recorded (Vmax $g^{-1}$ dw soil), F:B correspond to the fungal to bacterial ratio abundance measured in gene copy number with qPCR of ITS and 16 S rRNA gene (copy number $g^{-1}$ dw soil) and BG-intercept and NAG-intercept correspond to the enzymatic activity estimated at 0$^{o}$C (A); relationships between SOM NMDS axis 1 and dissolved organic carbon across warmed and control soils in mineral and organic horizons.

**Supplementary Table 2:** Metagenomes accession numbers obtained during this study generated at JGI.

| Sequencing Project ID | Library Name | Project Accession | Run Accession |
|---|---|---|---|
| 1282648 | HBPPC | SRP382445 | SRR19747141 |
| 1282649 | HBPPG | SRP382452 | SRR19747148 |
| 1282650 | HBPPH | SRP382446 | SRR19747142 |
| 1282651 | HBPPN | SRP382447 | SRR19747143 |
| 1282652 | HBPPO | SRP382448 | SRR19747144 |
| 1282653 | HBPPP | SRP382449 | SRR19747145 |
| 1282654 | HBPPS | SRP382450 | SRR19747147 |
| 1282655 | HBPPT | SRP382456 | SRR19747151 |
| 1282656 | HBPPU | SRP382451 | SRR19747146 |
| 1282657 | HBPPW | SRP382459 | SRR19747155 |
| 1282658 | HBPPX | SRP382453 | SRR19747149 |
| 1282659 | HBPPY | SRP382461 | SRR19747157 |
| 1282660 | HBPPZ | SRP382457 | SRR19747153 |
| 1282661 | HBPSA | SRP382458 | SRR19747154 |
| 1282662 | HBPSB | SRP382462 | SRR19747158 |
| 1282663 | HBPSC | SRP382454 | SRR19747150 |
| 1282664 | HBPSG | SRP382460 | SRR19747156 |
| 1282665 | HBPSH | SRP382464 | SRR19747160 |
| 1282666 | HBPSN | SRP382455 | SRR19747152 |
| 1282667 | HBPSO | SRP382463 | SRR19747159 |
